# Cholecystokinin exerts a major control on corticostriatal synapse and motor behavior

**DOI:** 10.1101/2022.11.28.518143

**Authors:** Chloé Guillaume, María Sáez, Blandine Castellano, Patricia Parnet, Ramon Reig, Vincent Paillé

## Abstract

Cholecystokinin (CCK) is a neuropeptide detected and produced at high concentrations in the brain. To date it was mainly used as a neuronal marker of neuronal subtypes and its role as a neuromodulator was poorly known. However, few studies showed that it could be an essential neuromodulator in various brain structures, playing a role on synaptic plasticity and memory consolidation. In order to better understand the processes by which CCK impacts plasticity, we focus our attention on the striatum, a nucleus involved in procedural learning and motor behavior, with a rich expression of CCK receptor type 2 (CCK2R). By using *in-vivo* and *ex-vivo* electrophysiological approaches, we show that CCK is involved in the corticostriatal synaptic transmission and has a key role in its plasticity. Using *in-vivo* optopatch-clamp of identified MSNs, we observe a decrease of corticostriatal synaptic transmission after an injection of CCK2R antagonist, leading to a reduction of evoked excitatory post synaptic potential recorded on both MSNs populations (direct and indirect pathways). In addition, we evaluate the impact of CCK2R antagonist on corticostriatal synaptic plasticity using Spike Timing Dependent Plasticity (STDP) protocols on MSNs of acute rat brain slices. Results demonstrate that the CCK2R antagonist is able to reverse the corticostriatal synaptic plasticity (i.e. LTP protocol leads to LTD). Finally, we evaluate the effect of CCK2R antagonist on the motor behavior of juvenile rats challenged with different locomotor tests and show a sex-dependent impairment of motor behavior. Overall, our results demonstrate that CCK and its receptor CCK2R are essential for inputs processing encoding in the corticostriatal network with consequences on motor activity.

**Significant statement:** Cholecystokinin (CCK) is considered to be one of the most abundant neuropeptides in the brain but its role as a neuromodulator is not well understood. In our study we investigate its role on the corticostriatal transmission which is a well characterized synapse highly involved in motor and cognitive functions. Here, we show that CCK2R is crucial for the corticostriatal synaptic transmission and plasticity. Indeed, CCK binding on CCK2R is essential for LTP induction by STDP. Finally, we demonstrate that the blockage of CCK2R affects corticostriatal synaptic transmission and motor ability in male rats.

## Introduction

Cholecystokinin (CCK) is considered as the hormone of satiety since its discovery in the gastrointestinal tract. Its role at the periphery, secreted by enteroendocrine cells, has been well characterized; however, since the 80s it is known that CCK is widely distributed in the central nervous system and is even considered as the most abundant neuropeptide in the brain (Moran and Schwartz, 1994). CCK is present in almost all brain regions but with particular high levels in the cortex and limbic regions (Vanderhaeghen et al., 1980; Beinfeld et al., 1981; Crawley, 1985). Its coexistence with several neurotransmitters, such as dopamine or GABA, suggests an important role in the neuromodulation (Somogyi et al., 1984; Crawley et al., 1985). Both forms of CCK specific receptors CCK-1 and CCK-2 (CCK1R and CCK2R respectively) are also found in the brain (Innis and Snyder, 1980; Hill et al., 1987). However, CCK2R is the major form in the brain (Van Dijk et al., 1984) and has a significantly high expression level in the cortex, nucleus accumbens (NAc), hypothalamus and amygdala, as well as in the striatum (Pélaprat et al., 1987). The role of CCK in the central nervous system is very poorly understood. In the past, studies have mainly focused on how CCK-positive interneurons contributed to the function of hippocampal and cortical circuits (Lee and Soltesz, 2011), but most studies have considered CCK in the brain only as a molecular marker of neuronal subtypes without evaluating its potential neuromodulator role. Nevertheless, an increasing number of studies have demonstrated the importance of CCK at different levels, from cellular to behavior (Lau et al., 2022). In rodents, in the hippocampus, CCK2R activation by CCK8s increases GABA release (Deng and Lei, 2006) and increases excitatory postsynaptic current (EPSC) by enhancing releasable glutamate vesicles (Deng et al., 2010). Previously, it was shown to mediate the release of dopamine in the NAc but to reduce the affinity of dopamine for D2 receptors (Ferraro et al., 1996). CCK8s is also involved in synaptic plasticity: in the hippocampus, by facilitating long term potentiation (LTP) (Balschun and Reymann, 1994; Yasui and Kawasaki, 1995) and in the dorsomedial nucleus of the hypothalamus, by shifting the GABA synapses from long term depression (LTD) to LTP (Crosby et al., 2018). Concerning its impact on the behavior, CCK induces a state of anxiety in rodents (Fekete et al., 1984; Biró et al., 1993; Daugé and Léna, 1998a; Sadeghi et al., 2017) but also in humans and primates (Bradwejn et al., 1992; Palmour et al., 1992; Harro et al., 1993). It is also worth mentioning that CCK can modulate learning and memory processes in rodents with memory-enhancing effects (Voits et al., 1995, 2001; Sebret et al., 1999; Hadjiivanova et al., 2003; Nguyen et al., 2020). In 1981, it was highlighted that in Huntington’s disease there is a diminution of 75% of the CCK receptors in the patients’ caudate-putamen, suggesting a possible implication of CCK in the impaired motor processes generally observed in this disease (Hays et al., 1981). Given the presence of CCK and CCK2R in high concentration in the cortex and in the striatum and the presence of CCK in the corticostriatal glutamatergic synapse (Hökfelt et al., 2002), it is essential to understand the role of CCK in corticostriatal transmission and plasticity.

The corticostriatal pathway is responsible for transferring and filtering cortical information to the basal ganglia, the striatum being the entry point (Smith and Bolam, 1990), and its plasticity is considered to be one of the cellular substrates absolutely crucial for procedural learning (Wickens, 2009). In fact, the synaptic plasticity in the striatum is central for several striatal functions such as memory formation of sensorimotor information and motor control (Calabresi et al., 1996; Kreitzer and Malenka, 2008; Di Filippo et al., 2009). Furthermore, that plasticity is often altered in several neuropathologies affecting basal ganglia (Kreitzer and Malenka, 2008; Calabresi et al., 2009).

To decipher the importance of CCK in the corticostriatal transmission, we performed *in-vivo* and *ex-vivo* electrophysiological and behavioral tests involving young rats. We show for the first time that CCK plays a critical role in corticostriatal synaptic plasticity. Indeed, CCK2R blockage (using LY225910) induces a switch of the corticostriatal plasticity by leading to an LTD in place of LTP. CCK also plays a key role in the corticostriatal synaptic transmission since CCK2R blockage induces a diminution of evoked EPSP and EPSC in MSNs following a cortical stimulation *in-vivo* and *ex-vivo*. Finally, we are able to show that *in-vivo* blockage of CCK2R impairs motor coordination in rats.

## Materials and Methods

### Materials

*In-vivo* electrophysiology (Table 1)

**Table 1:** *In-vivo* electrophysiology material.

|  | *Product* | *Provider* | *Features* | *Comments* |
| --- | --- | --- | --- | --- |
| **Animals** | **Rats Sprague-Dawley** | Janvier Lab, Le-Genest-Saint-Isles, France | Postnatal day 14 to 35 |  |
| **Drugs** | **LY225910 (LY)** | Tocris, Ellisville, Missouri, USA | N° lot: 2B/235860 | CCK2R antagonist ; use at concentration 1 μM |
| | **Lorglumide** | Sigma-Aldrich, St-Quentin, France | Serie number: BCBW5189 | CCK1R antagonist ; use at concentration 1 μM |
| **Solutions** | **Extracellular solution** | Made at lab (Chemicals are purchased from Sigma, St Quentin, France) | NaCl 125 mM, KCl 2.5 mM, CaCl<sub>2</sub> 2 mM, MgCl<sub>2</sub> 1 mM, NaHCO<sub>3</sub> 25 mM, Glucose 25 mM, NaH<sub>2</sub>PO<sub>4</sub> 1.25 mM, 95% O<sub>2</sub>/5% CO<sub>2</sub> |  |
|  | **Intracellular solution** | Made at lab (Chemicals are purchased from Sigma, St Quentin, France) | Kgluconate 122 mM, KCl 13 mM, Hepes 10 mM, MgATP 4 mM, NaGTP 0.3 mM, Phosphocreatine 10 mM, pH 7.36 |  |
| **Equipment** | **Vibratome** | LEICA Microsystem, Germany | Leica VT 1000S |  |
|  | **Pump Minipuls 3** | GILSON INC, Middleton, Wisconsin, USA |  |  |
|  | **Borosilicate glass pipette** | Harvard apparatus, USA | Glass capillaries GC150TF-10 |  |
|  | **Pipette puller** | Narishige, Japan | PC-10 Dual stage Micropipette puller |  |
|  | **Microscope** | Olympus Corporation, Japan | Olympus BX51WIF |  |
|  | **Water-immersion objective X40** | Olympus Corporation, Japan | Olympus U-CAMD3 |  |
|  | **Bipolar electrode** | FHC Inc, Bowdoin, Maine, USA | CBDSF75 N°lot: 229732 |  |
|  | **Amplifier HEKA** | HEKA Instrument Inc, New-York, USA | HEKA EPC10 USB double |  |
| **Software** | **Patchmaster** | HEKA Instrument Inc, New-York, USA |  |  |
|  | **Matlab** | Mathworks, Natick, Massachusetts, USA | R2022a |  |
|  | **GraphPad Prism** | GraphPad Software, San Diego, California, USA | GraphPad Prism 9 |  |

*Ex-vivo* electrophysiology (Table 2)

**Table 2:**
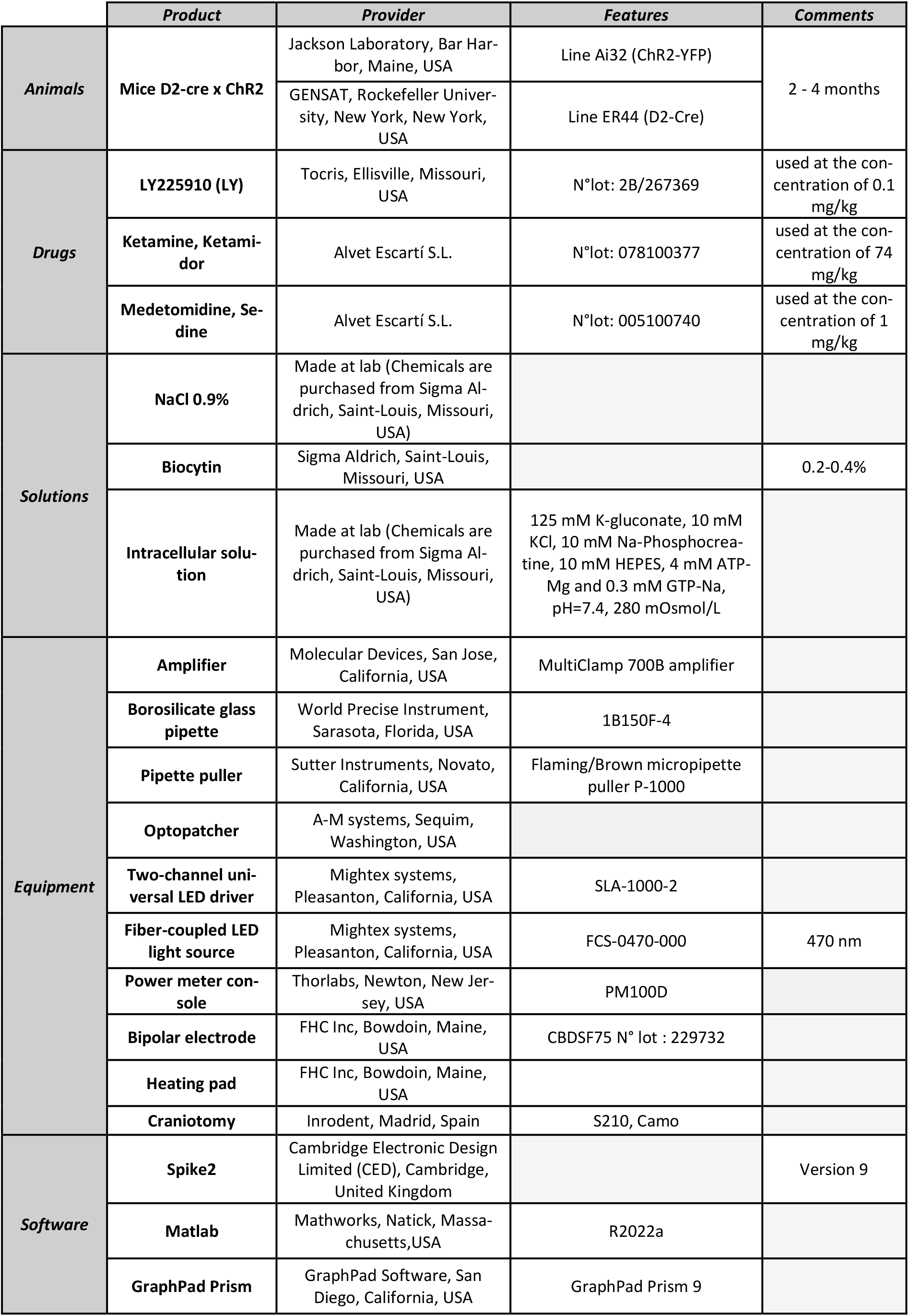
*Ex-vivo* electrophysiology material.

|  | *Product* | *Provider* | *Features* | *Comments* |
| --- | --- | --- | --- | --- |
| **Animals** | **Mice D2-cre x Chr2** | Jackson Laboratory, Bar Harbor, Maine, USA | Line Ai32 (Chr2-YFP) | 2 - 4 months |
|  |  | GENSAT, Rockefeller University, New York, New York, USA | Line ER44 (D2-Cre) |  |
| **Drugs** | **LY225910 (LY)** | Tocris, Ellisville, Missouri, USA | N°lot: 2B/267369 | used at the concentration of 0.1 mg/kg |
|  | **Ketamine, Ketamidol** | Alvet Escartí S.L. | N°lot: 078100377 | used at the concentration of 74 mg/kg |
|  | **Medetomidine, Sedine** | Alvet Escartí S.L. | N°lot: 005100740 | used at the concentration of 1 mg/kg |
| **Solutions** | **NaCl 0.9%** | Made at lab (Chemicals are purchased from Sigma Aldrich, Saint-Louis, Missouri, USA) |  |  |
|  | **Biocytin** | Sigma Aldrich, Saint-Louis, Missouri, USA |  | 0.2-0.4% |
|  | **Intracellular solution** | Made at lab (Chemicals are purchased from Sigma Aldrich, Saint-Louis, Missouri, USA) | 125 mM K-gluconate, 10 mM KCl, 10 mM Na-Phosphocreatine, 10 mM HEPES, 4 mM ATP-Mg and 0.3 mM GTP-Na, pH=7.4, 280 mOsmol/L |  |
| **Equipment** | **Amplifier** | Molecular Devices, San Jose, California, USA | MultiClamp 700B amplifier |  |
|  | **Borosilicate glass pipette** | World Precise Instrument, Sarasota, Florida, USA | 1B150F-4 |  |
|  | **Pipette puller** | Sutter Instruments, Novato, California, USA | Flaming/Brown micropipette puller P-1000 |  |
|  | **Optopatcher** | A-M systems, Sequim, Washington, USA |  |  |
|  | **Two-channel universal LED driver** | Mightex systems, Pleasanton, California, USA | SLA-1000-2 |  |
|  | **Fiber-coupled LED light source** | Mightex systems, Pleasanton, California, USA | FCS-0470-000 | 470 nm |
|  | **Power meter console** | Thorlabs, Newton, New Jersey, USA | PM100D |  |
|  | **Bipolar electrode** | FHC Inc, Bowdoin, Maine, USA | CBDSF75 N° lot : 229732 |  |
|  | **Heating pad** | FHC Inc, Bowdoin, Maine, USA |  |  |
|  | **Craniotomy** | Inrodent, Madrid, Spain | S210, Camo |  |
| **Software** | **Spike2** | Cambridge Electronic Design Limited (CED), Cambridge, United Kingdom |  | Version 9 |
|  | **Matlab** | Mathworks, Natick, Massachusetts, USA | R2022a |  |
|  | **GraphPad Prism** | GraphPad Software, San Diego, California, USA | GraphPad Prism 9 |  |

Behavioral tests (Table 3)

**Table 3:** Behavioral tests material.

|  | *Product* | *Provider* | *Features* | *Comments* |
| --- | --- | --- | --- | --- |
| **Animals** | **Rats Sprague-Dawley** | Janvier Lab, Le-Genest-Saint-Isles, France | Postnatal day 28 to 35 | housed in an inverted light/dark cycle (light off at 7 AM and light on at 7 PM), all behavioral tests are realized between 8 AM and 11 AM |
| **Drugs** | **LY225910 (LY)** | Tocris, Ellisville, Missouri, USA | N° lot : 2B/239941 | used at the concentration of 0.1 mg/kg |
|  | **CCK8** | Bachem, Bubendorf, Switzerland | N° lot : 1066802 | used at the concentration of 0,01 mg/kg |
|  | **NaCl 0,9%** | B. Braun Medical, Saint-Cloud, France |  |  |
| **Equipment** | **Rotarod** | Bioseb, Vitrolles, France |  |  |
| **Software** | **Video tracking system** | Videotrack software, Viewpoint, Lyon, France |  |  |
|  | **BORIS software** |  | Copyright 2012-2021 Olivier Friard - Marco Gamba |  |
|  | **GraphPad Prism** | GraphPad Software, San Diego, California, USA | GraphPad Prism 9 |  |

C-Fos Immunolabelling detection (Table 4)

**Table 4:**
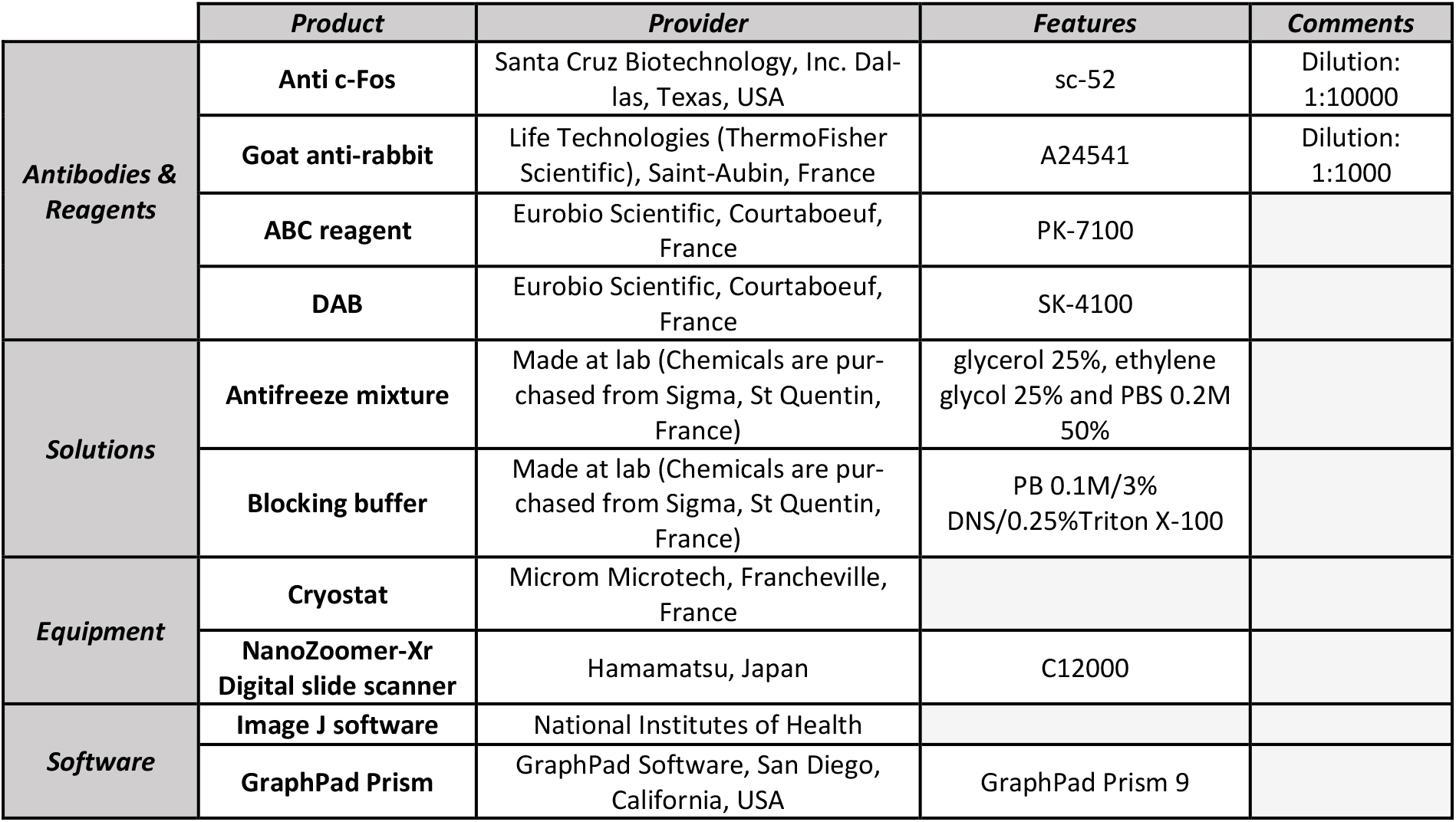
c-Fos Immunolabelling detection material.

Morphological reconstruction and BDA axonal tracing (Table 5)

**Table 5:** Morphological reconstruction and BDA axonal tracing material.

|  | *Product* | *Provider* | *Features* | *Comments* |
| --- | --- | --- | --- | --- |
| **Antibodies & Reagents** | **Cy3-conjugated streptavidin** | Jackson Immunoresearch laboratories Cambridgeshire, UK |  | Dilution: 1:1000 |
|  | **Biotinylated Dextran Amine** | Sigma Aldrich, Saint-Louis, Missouri, USA |  | 300 nl |
| **Solutions** | **Mowiol** | Calbiochem, San Diego, California, USA |  |  |
| **Equipment** | **Cryotome** | Leica, Wetzlar, Germany |  |  |
|  | **Stereotaxic** | Kopf instruments, Tujunga, California, USA |  |  |
|  | **Thunder Imager microscope** | Leica, Wetzlar, Germany |  |  |
| **Software** | **Image J software** | National Institutes of Health |  |  |
|  | **Leica Application Suite X software** | Leica, Wetzlar, Germany |  |  |

### Methods

#### Animals

##### Rats

All rats were obtained from Janvier Lab and all experimental protocols were performed in accordance to the local ethical committee on animal experimentation (CEEA PDL) and the approval from the French Ministry of Superior Education and Research (MESR) (APAFIS #30245-202104291141254v1).

*Ex-vivo* electrophysiology experiments were performed using young Sprague-Dawley rats (postnatal day 14 to 35). Intra-peritoneal (i.p.) injection of 0.1 ml of pentobarbital is used to anaesthetize the animal 10 min before decapitation.

Behavioral motor tests were performed using young Sprague-Dawley rats (postnatal day 28 to 35). Animals were housed in an inverted light/dark cycle (light off at 7 AM and light on at 7 PM), and all behavioral tests were performed between 8 AM and 11 AM.

##### Mice

All the experimental procedures performed on mice were conformed to the directive 2010/63/EU of the European Parliament and of the European Council, and the RD 53/2013 Spanish regulation on the protection of animal use for scientific purposes, approved by the government of the Autonomous Community of Valencia, under the supervision of the Consejo Superior de Investigaciones Científicas and the Miguel Hernandez University Committee for Animal use in Laboratory.

*In-vivo* electrophysiology experiments were performed using transgenic mice D2-cre × ChR2 of both sexes (n= 23; n= 12 females and 11 males) between 2-4 months of age in order to induce the expression of channelrhodopsin for the identification of direct and indirect MSNs as previously shown (Alegre-Cortés et al., 2021; Ketzef et al., 2017). To obtain this transgenic line, a D2-cre (ER44 line, GENSAT) mouse line was crossed with the channelrhodopsin (ChR2)-YFP reporter mouse line (Ai32, The Jackson Laboratory). To perform *in-vivo* electrophysiology recordings, mice were anesthetized by an intraperitoneal injection of ketamine (74 mg/kg) and medetomidine (1 mg/kg) diluted in 0.9 % NaCl. A maintaining dose of ketamine (30 mg/kg i.p.) was administered every 2 hours or after changes in the EEG or reflex responses to paw pinches. After experimentation, animals were sacrificed by receiving an overdose of sodium pentobarbital (200 mg/kg i.p.).

#### Drugs

For *in-vivo* electrophysiology experiments, only LY was used on mice. LY previously dissolved in DMSO was used at the concentration 0.1 mg/kg. LY was administered i.p. to the mice while performing whole-cell patch clamp recordings and the analysis of any effect on the spontaneous activities or in the PSTH evoked potential was conducted at least 15 minutes post-injection.

For *ex-vivo* electrophysiology experiments, three different drugs were used on rats’ horizontal brain slices obtained after brain extraction and slicing with a vibratome (described below). LY225910 (LY) is a CCK2R antagonist (IC_50_ = 9.3 nM). After its dissolution in DMSO, LY solution was added to the extra-cellular solution to obtain the perfusion solution of LY at 1μM (Crosby et al., 2018). Lorglumide is a CCK1R antagonist. Lorglumide solution was added to the extracellular solution to obtain the perfusion solution of 1μM (Crosby et al., 2018).

For behavioral tests, two different drugs were used. LY previously dissolved in DMSO was used at the concentration 0.1 mg/kg (following (Farook et al., 2004). To reduce bias between the LY and the sham groups, the sham group received a solution of NaCl 0.9% with 0,4% of DMSO. CCK8 was used at the concentration 0.01 mg/kg. All injections were i.p..

#### Electrophysiology

##### *In-vivo* recordings

Tracheotomy was performed to increase mechanical stability during recordings by decreasing breathing-related movements. Mice were placed in a stereotaxic frame (customized Stoelting stereotaxic base) and air enriched with oxygen was delivered through a thin tube placed 1 cm from the tracheal cannula. Temperature was maintained at 36.5 ± 0.5°C using a feedback-controlled heating pad. Three craniotomies were drilled at different sites from bregma: AP 0 mm, LM 4 mm (Dorsolateral striatum, whole-cell recordings); AP −0.7 mm, LM 3.8 mm (ipsi S2, stimulation); AP −0.7 mm, LM −3.8 mm (contra S2, LFP recordings) (following *Paxinos and Franklin 2001*).

Whole-cell patch-clamp recordings were obtained from the dorsolateral striatum between 2,068 and 2,251 μm deep at a penetration angle of ^~^30°. The exposed brain was continuously covered by 0.9% NaCl to prevent drying. Signals were amplified using a MultiClamp 700B amplifier and digitized at 20 KHz with a CED acquisition board and Spike 2 software. Borosilicate patch pipettes were pulled with a Flaming/Brown micropipette puller P-1000 and had an initial resistance of 6-12 MΩ, with extended tips to minimize cortical damage. Pipettes were back-filled with intracellular solution containing: 125 mM K-gluconate, 10 mM KCl, 10 mM Na-Phosphocreatine, 10 mM HEPES, 4 mM ATP-Mg and 0.3 mM GTP-Na. pH and osmolarity were adjusted to ^~^7.4 and ^~^280 mOsm/L, respectively. Biocitin was then added to the intracellular solution to reconstruct the recorded cell after the experiment. For all neurons, the input resistance was measured as the slope of a linear fit between the injected current steps and membrane potential of 2 s duration. The time constant (tau) was calculated as the time required for the membrane voltage change to reach 63% of its maximum value. Capacitance was obtained by dividing the time constant between the voltage membrane resistance. The average recording time for all MSNs was 68.6 ±34.95 min (minimum=42 min, maximum=156 min; n=23). Recorded MSNs were optogenetically identified and classified as a putative direct or indirect MSN by means of the optopatcher (Katz et al., 2013). In order to identify the specific type of MSNs belonging to the direct and indirect pathways, the optopatcher was used (Katz et al., 2013, 2019; Ketzef et al., 2017a; Alegre-Cortés et al., 2021) (A-M systems). Pulses (Two-channel universal LED driver) of blue light (Fiber-coupled LED light source) controlled through Spike 2 software were delivered using an optical fiber (200 μm diameter, handmade) inserted into the patch-pipette, while recording their spontaneous activity. One or two serial pulses with 5 light steps of 500 ms each were delivered every 2 seconds with increasing intensity from 20% to 100 % of full LED power (minimal light power of 0.166 mW; maximal power of 0.83 mW at the tip of the fiber). The light power was measured with a power meter console. Positive cells responded to light pulses by depolarizing their membrane potential, whereas negative cells did not show any depolarization to light pulses. We then studied their electrophysiological properties by injecting positive and negative intracellular current steps. Finally, electrical stimulation was performed in S2 with a bipolar electrode. The stimulation consisted of a pulse of 10 ms duration and 6.5 μA intensity.

##### *Ex-vivo* recordings

We used the same brain slice orientation than previously described (Fino et al., 2005; Paille et al., 2013). After extraction, the brain was placed in a frozen oxygenated extracellular solution. Horizontal slices of 300 μm thickness were obtained with vibratome to preserve intact connections between cortex and striatum. Slices were then placed in an oxygenated extracellular solution at 32°C for one hour. In the electrophysiology station, slices were continuously perfused with extracellular solution at room temperature with a peristaltic pump (Pump Minipuls 3). A borosilicate glass pipette filled with intracellular solution was placed on the Medium Spiny Neuron (MSN) in whole-cell patch-clamp configuration. Individual neurons were imaged through a microscope with a X40 water-immersion objective. For the stimulation protocol, a bipolar electrode was placed in the V layer of the somatosensory cortex. The cortical stimulations were monophasic and at constant current between 3 and 32 mA with a duration range of 100 – 150 μs. Data were recorded using a HEKA amplifier and acquisition software Patchmaster. As previously described (Paillé, Fino 2013), the 4 recorded steps are: Step 1) The baseline measurement consisted on recording the MSN excitatory post-synaptic current (EPSCs) until a regular response of MSN to the cortical stimulation was obtained (20 min). Step 2) After the baseline, the pharmacological solution (LY or Lorglumide) was added into the bath to record MSN EPSC in presence of drugs thus constituting the baseline + drug step. For the control group, recordings were performed without drug perfusion. Step 3) STDP protocol was performed by pairing pre- and postsynaptic stimulations repeated 100 times at 1 Hz with a time shift of a few milliseconds (Δt). The presynaptic stimulation corresponds to the cortical stimulation and the postsynaptic stimulation corresponds to the depolarization of the MSN leading to an evoked AP. Step 4) MSN EPSCs were recorded for 60 min. MSN EPSCs amplitudes were compared to EPSCs amplitudes obtained during step 2 to assess the occurrence of long-term synaptic plasticity changes. Signals were digitized at 10 kHz. Recordings were corrected for a junction potential of +18 mV. The series resistance was monitored throughout the experiment by a brief voltage step of −5 mV at the end of each recording. Data were discarded when the series resistance increased or decreased by >20% compared to the baseline. Data analysis of the recordings was performed with MatLab.

#### Behavioral test

The rats were randomly assigned to three groups in order to be injected with NaCl 0,9% for the sham group, LY225910 solution for the LY group, CCK8 solution for the CCK group. Rats were weaned at 21 postnatal days and started the habituation phase: two experimenters gently manipulated each rat for 2 min every day for 5 days to allow the animal familiarizing with the experimenters and the behavioral assessment equipment. This manipulation took place in the experimental room with the Rotarod running so that the rats became familiar with the room, the transportation in the room, and the noise of the Rotarod in operation. The timeline of the behavioral study is shown in Figure 4A.

**Figure 1:**
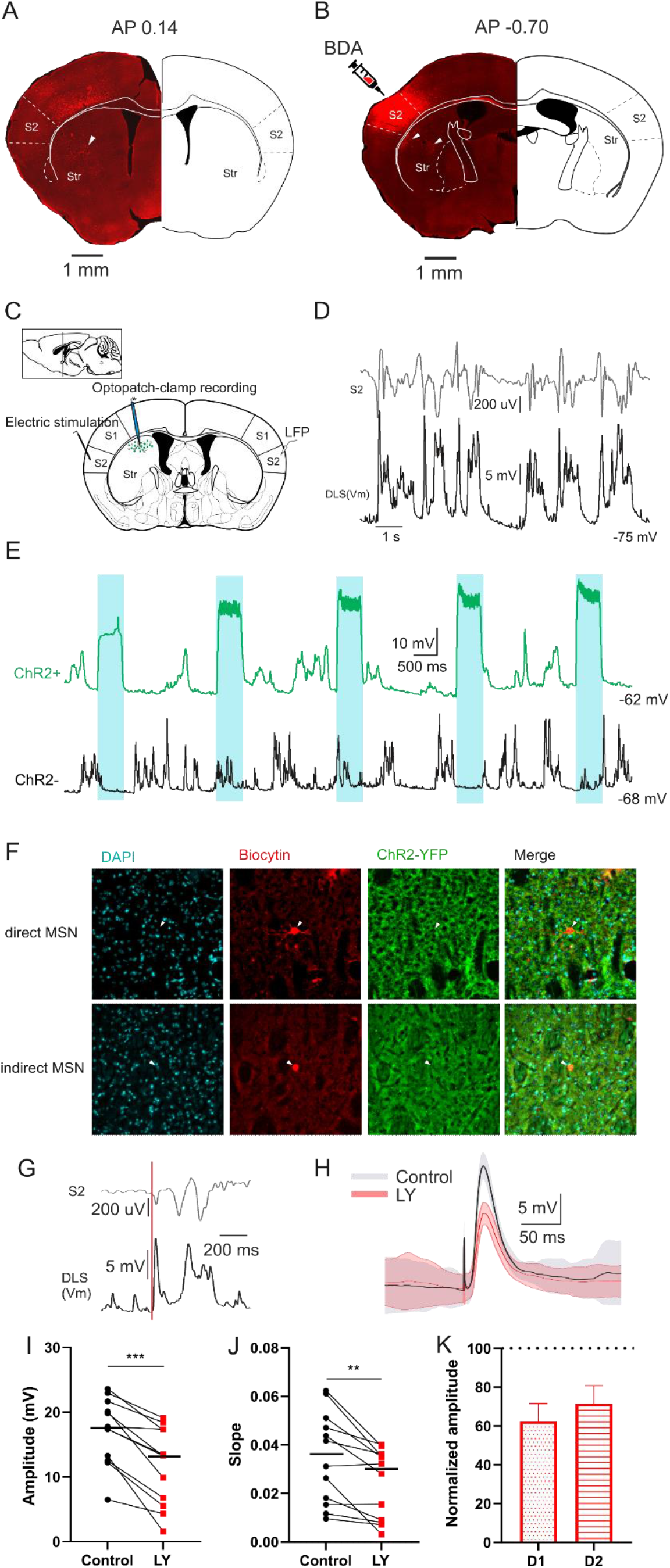
LY injection decreases evoked glutamatergic response *in-vivo* on both direct and indirect pathways MSNs. **A** Coronal section with projections in the striatum from the injection site in S2. **B** Coronal section with injection site in S2 and projections in the striatum. **C** Stimulation electrode in S2, optopatch pipette in dorsal striatum and LFP electrode in contralateral S2. **D** Representative recorded traces. **E** Discrimination of D1 (ChR2-) or D2 (ChR2+) MSN by pulsed of blue light. **F** Morphological reconstruction of direct MSN (D1) above and indirect MSN (D2) below. **G** Representative trace of the records in dorsolateral striatum (DLS) and S2 contralateral, red line is the stimulation artefact. **H** Average trace of the post-synaptic potential (PSP) in downstate before and after injection of LY. **I** Significant diminution of the PSP amplitude in LY condition (***p=0.0005, n=12, Wil-coxon signed rank test). **J** Significant diminution of the EPSP slope in LY condition (**p=0.0024, n=12, Wil-coxon signed rank test). K PSP normalized amplitude after LY injection do not presents significant differences between both D1 MSNs (n=8) and D2 MSNs (n=4) (respectively direct and indirect path-ways).

**Figure 2:**
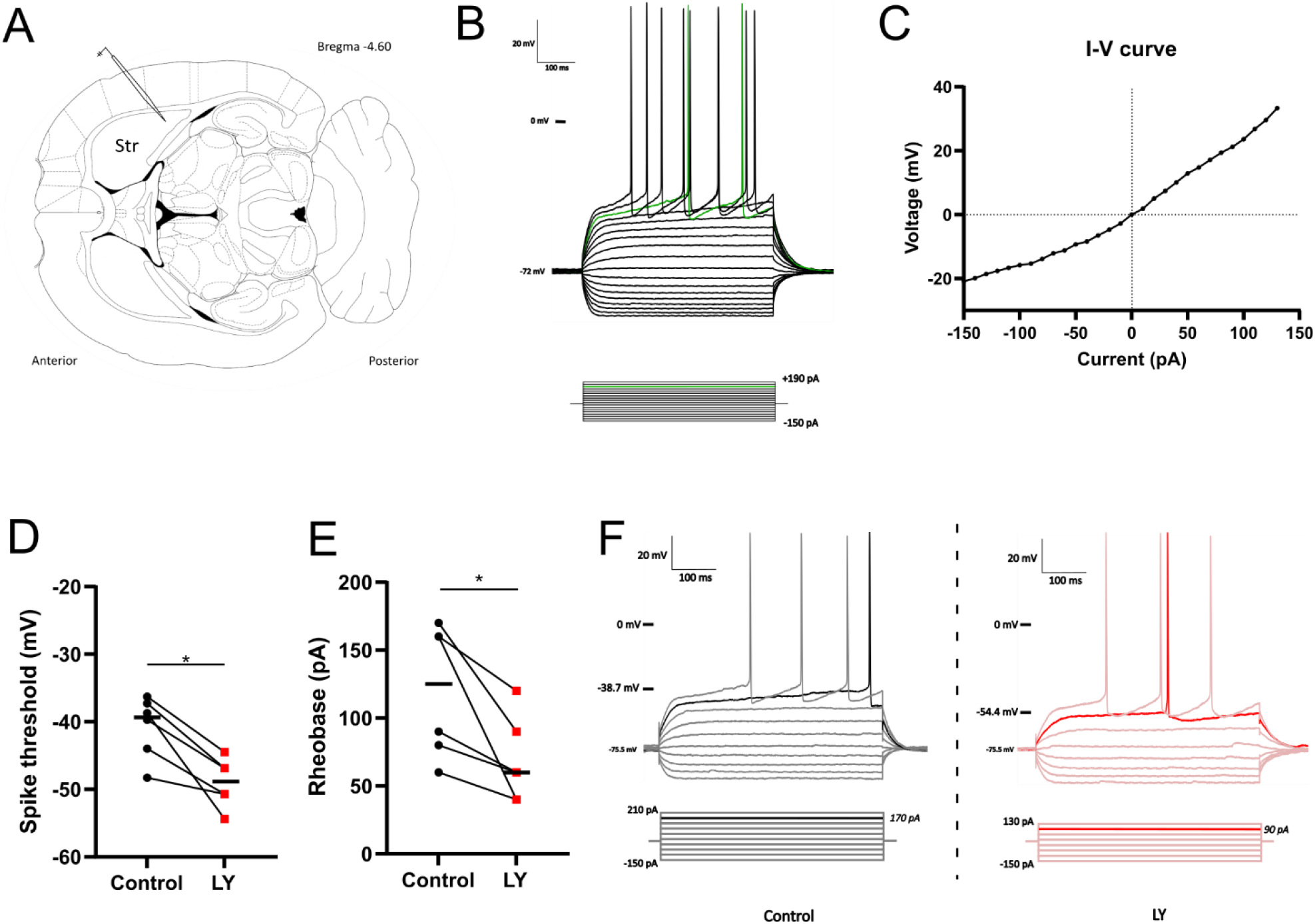
Recorded MSNs characteristics and LY effect on membrane actives properties. **A** Patch-clamp pipette on the recording area. **B** Below: current injection protocol, current applied for 500 ms, first at −150 pA, then increment by 10 pA every 4 seconds (not each steps are shown), rheobase is in green. Above: MSN response to the current injection protocol, hyperpolarisation, depolarisation, first firing (green) and spiking pattern. **C** I-V curve from a representative recorded MSN. **D** Significant diminution of spike threshold after LY infusion (*p=0.0313, n=6, Wilcoxon signed rank test). **E** Significant diminution of rheobase after LY infusion (*p=0.0313, n=6, Wilcoxon signed rank test). **F** Spike recorded traces from a representative recorded MSN before and after LY infusion.

**Figure 3:**
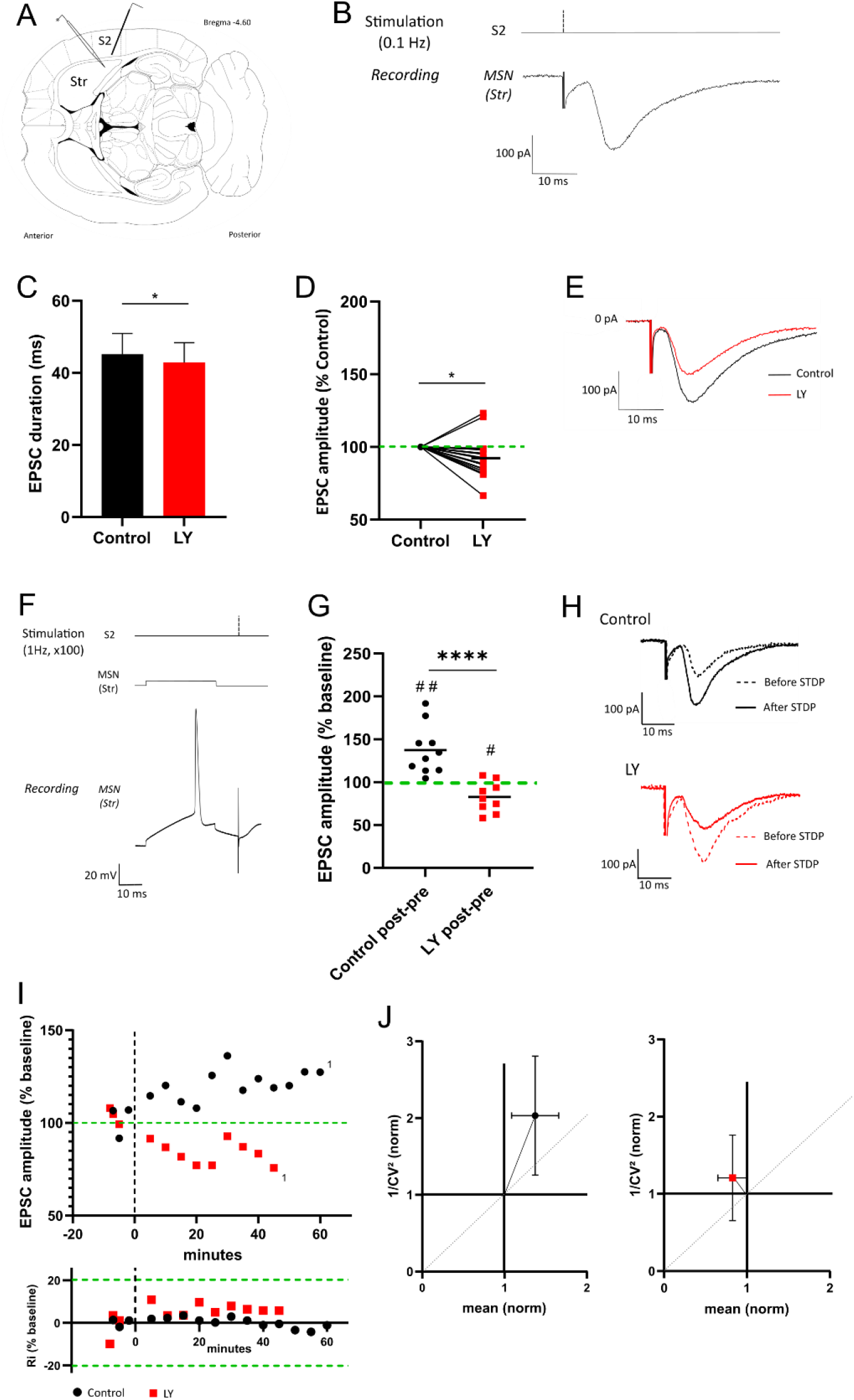
LY affects the evoked glutamatergic response and switch the STDP long term potentiation *ex-vivo*. **A** Stimulation electrode placed in S2 and a patch clamp pipette in the recording area (dorsolateral part of the striatum). **B** Stimulation protocol, repeated every 10 sec i.e. 0.1 Hz in S2 and MSN recorded response. **C** Significant EPSC duration decrease after LY infusion (*p=0.0237, n=18, Wilcoxon signed rank test). **D** EPSC amplitude in LY condition in % of the control condition (*p=0.0208, n=18, Wilcoxon signed rank test). **E** Evoked glutamatergic response from a representative recorded MSN in control and LY condition. **F** STDP protocol, post-pre pairing 100 times at 1Hz. **G** Normalised EPSC amplitude in control condition and in LY condition, (control n=10 and LY n=9, ****p<0.0001 control vs LY, Mann-Whitney test, #p=0.0190 LY vs baseline, one sample t test, ##p=0.0026 control vs baseline, one sample t test). **H** EPSC from representative MSN in each condition during baseline (before STDP) and long term (after STDP). **I** Time monitoring of EPSC amplitude and resistance for representative MSN in control and in LY condition. **J** Normalised 1/CV^2^ of control condition and LY condition.

**Figure 4:**
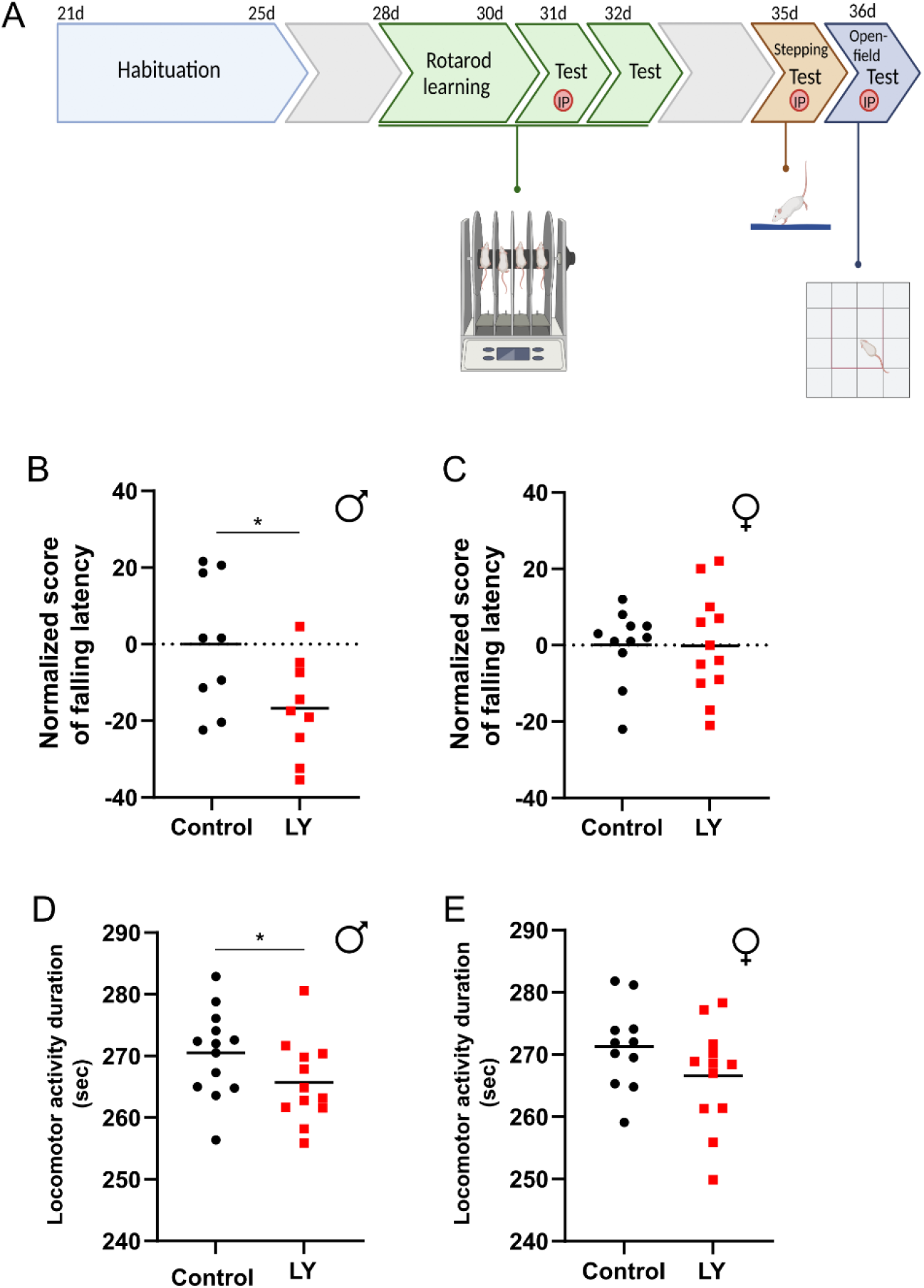
LY affect males motor behavior. **A** Time-line of the behavioral study. (5 days of habituation, Rotarod learning from 28 to 30 days, Rotarod test at 31 days after an i.p. injection, the Rotarod test at 32 days without injection, Stepping-test. at 35 days after an i.p. injection, Open-field test at 36 days after an i.p. injection. **B** Rotarod normalized score of falling latency (difference of falling latency between with and without injection of NaCl or LY, normalized to the value of the control group) for males’ rats (*p= 0.0331, n=12, unpaired t test). **C** Rotarod normalized score of falling latency for females’ rats in the Rotarod (control n=11, LY n=12). **D** Male rats locomotor activity duration in the Open-field (*p=0.0457, control n=13 and LY n=12, Mann-Whitney). **E** Females rats locomotor activity duration in the Open-field (control n=11 and LY n=12).

##### Rotarod performance test

The rotarod performance test is based on a rotating rod with forced motor activity being applied. The test is evaluating balance, grip strength and motor coordination of the subjects. We used 28 days aged rats. For the acquisition phase, the rat was placed on the rotating cylinder at a fixed speed of 15 rpm. If the rat remained on the cylinder for at least 1 min, the task was considered acquired and the test was stopped. To learn the task, the rat was trained every day for 3 days and performed 3 trials with 5 min rest intervals in their housing cage. At 31 postnatal days, rats performed the test 30 min after receiving an i.p. injection of the drug (Sham, LY or CCK). In the test condition, the cylinder is accelerated up to 40 rpm after one minute. At 32 postnatal days, the rat performed the same test without drug injection in order to compare its performance with and without injection. The rat’s performance was assessed by the time it stood on the cylinder in the first trial.

##### Stepping test

This test assesses the rat’s postural recovery and is performed on a table covered with a mat to prevent the animal from slipping on a smooth surface. The rat was placed at one end of the table with all four paws touching the surface. The experimenter held the rat by the hind limbs with one hand, slightly lifting the lower body of the animal above the surface. The animal was dragged across the surface (covering a distance of 1 meter) in 3-4 seconds.

The movement was performed three times backwards and three times forwards. There was no training session. At 35 postnatal days, the rats performed the test 30 min after having received the drug corresponding to their group (Sham, LY, CCK). The test was filmed and each video was then analyzed with BORIS. For each video the following parameters were recorded: number of postural recoveries with the right front paw, number of postural recoveries with the left front paw, number of vocalizations, time spent dragging on the mat, time spent turning.

##### Open-field test

The Open-field (OF) test is an experimental test used to address general locomotor activity levels, anxiety, and willingness to explore. Each arena of the Open-field apparatus measured 50 cm × 50 cm and were made of white opaque plastic wall. At 36 postnatal days, rats performed the test 30 min after receiving the drug corresponding to their group (Sham, LY, CCK). The rats were placed in the center of the arena. Their movements were recorded for 5 min using the infra-red video tracking system Viewpoint. From these 5 minutes of free moving, we extracted different information using the videotrack software. First, we evaluated the activity time and the total travelled distance. Then we assessed the type of travel (small or large distances). To finish, we evaluated the entries and the time spent in the central area which is often use to qualify the level of anxiety of the animal.

#### Histology

##### C-Fos Immunolabeling detection

At 38 postnatal days, half of the rats were injected with the drug corresponding to their group. 90 min after drug injection rats were anaesthetized with isoflurane and perfused with a transcardial physiological saline perfusion followed by ice-cold 4% paraformaldehyde in phosphate buffer (PB, pH 7.4). The brain was rapidly removed, immersed in the same fixative for 10 day at 4°C, and finally stored in 30% PB sucrose for 48-72h. The brain was then frozen in isopentane at around −60°C, and finally stored at −80°C until use. The striatum was cut into 40 μm free floating serial coronal sections with a cryostat. We used slices from Bregma 2.20 mm to Bregma −0.80 mm. Free-floating slices were stored at −20°C in an antifreeze mixture (glycerol 25%, ethylene glycol 25% and PBS 0.2M 50%) until their utilization. Wells containing serial striatum-floating sections were first rinsed in PB 0.1M to eliminate anti-freeze solution. Then endogenous peroxidase was deactivated by incubating slices in peroxidase block (0.3% H_2_O_2_ solution) for 30 min with gentle agitation. Slices were rinsed in PB 0.1M and then incubated 2h in blocking buffer (PB 0.1M/3% DNS/0.25%Triton X-100). Free-Floating slices were incubated for 48 hours at 4°C with the primary antibody anti-c-Fos (1 :10000) in a blocking buffer. After incubation with primary antibody and subsequent washing in PB 0.1M, free-floating slices were incubated with second antibody: goat anti rabbit biotin (1 :1000) for 2h. Sections were rinsed and then incubated with ABC kit for 30 min. After rinsing, sections were placed in solution for DAB reaction with Nickel. Once the background was high enough the reaction was stopped by placing sections into water. Then sections were air-dried for 2 days and mounted in a dehydrated medium. For each rat, c-Fos-positive cells were counted at three different rostrocaudal levels of the striatum. For the left and the right side, a digitized picture comprising the whole striatum was obtained using X20 magnification of a NanoZoomer-Xr Digital slide scanner. To obtain the least biased estimate of the total number of neurons, we used the dissector principle and random systematic sampling (Sterio, 1984; Coggeshall, 1992). Lines were drawn around the dorsolateral, dorsomedial, ventrolateral and ventromedial parts of the striatum for each section. Using Image J software (cell counter plugin), c-Fos immunoreactive body cells were counted based on a stereological-like technique (unbiased quantization technique) to limit the bias of “over-counting”. A counting grid was randomly positioned on the image (“systematic random”), each small square having a side length of 100 μm and only the c-Fos positive cells present in one square out of four were counted by an observer blind to the rat number. Once the counts were concluded, the total density of the area was extrapolated according to the area of the analyzed structure. Data were expressed as the mean of c-Fos positive cells/mm^2^.

##### Morphological reconstruction of MSN patched *in-vivo*

At the end of each *in-vivo* experiment, mice were sacrificed with a lethal dose of sodium pentobarbital and perfused with a solution containing 4% paraformaldehyde in 0.1 M phosphate buffer (PB, pH 7.4). Brains were extracted and stored in a PBS solution until the cutting. Before cutting, brains were transferred into PBS containing 30% sucrose for at least 48 hours. Coronal slices (20 μm thick) containing the entire striatum from the recorded side (from AP 1.4 mm to AP −1.3 mm, following Paxinos and Franklin, 2001), were obtained using a digital automatic cryotome and collected on gelatin coated slides. Sections were incubated overnight with Cy3-conjugated streptavidin diluted (1:1000) in 1% BSA, 0.03% Triton-X 100 in 0.1 M PBS. Finally, the glass slides were covered with mowiol and imaged. Neurons were then reconstructed using the widefield Thunder Imager microscope and then the images were processed using ImageJ software.

##### Biotinylated Dextran Amine (BDA) axonal tracing in mice

In order to trace the projections from S2 towards the striatum, C57BL/6J mice (n=4) were anaesthetized with isoflurane and immobilized in a stereotaxic frame. 300 nl of BDA were injected unilaterally in S2 (at the following coordinates: AP −0.7 mm, LM 3.8 mm, DV −0.8 mm). For the anatomical study, mice were sacrificed 10 days after the injection by receiving an overdose of sodium pentobarbital (200 mg/kg i.p.). Brains were then extracted and processed as described above and then coronal slices (50 μm thick) containing both hemispheres from both sides were obtained using a digital automatic cryotome and collected on gelatin coated slides. Sections were incubated overnight with Cy3-conjugated streptavidin diluted (1:1000) in 1% BSA, 0.03% Triton-X 100 in 0.1 M PBS. Finally, the glass slides were covered with mowiol and imaged. Images were visualized, obtained and processed with the widefield Thunder Imager microscope and the Leica Application Suite X software.

#### Experimental design and statistical tests

For electrophysiology, groups data consisted of neuronal recordings from animals of both sexes. For *ex-vivo* electrophysiology, statistic evaluation of the effect of the drug on EPSC is performed on baseline measurements before drug infusion (the previous 20 minutes) and on baseline measurements after drug infusion (the next 20 minutes). Statistic evaluation of the occurrence of long-term plasticity is performed on the baseline measurement before STDP (the previous 20 minutes) and the longest moment where the cell is still alive (5 minutes selected from 30 minutes to 60 minutes after STDP). For the *in-vivo* electrophysiology, statistic evaluation of the effect of the drug on EPSP is performed on baseline measurements before injection (for 10 minutes) and on measurements 25 min after drug injection (for 10 minutes). For the behavior tests, the experimental design is described in Figure 4A. Group data are shown by (i) scatter plot of individual values with median bar for appaired data, (ii) scatter plot of individual values with mean bar for Rotarod test results, (iii) mean ± SEM (standard error of the mean), (iv) box and whisker plots for Kruskal-Wallis estimation. All statistical analyses are conducted with GraphPad Prism 9. Data are first tested with a normality test before performing appropriate statistical analyses including Wilcoxon signed rank test, Mann-Whitney test, one sample t test, unpaired t test and Kruskal-Walis. Statistical significance was defined as p<0.05.

## Results

### CCK2R inhibition decreases evoked glutamatergic response *in-vivo* on both direct and indirect pathway MSNs

To check the projections innervating dorsal striatum from S2 on mice, we injected BDA in S2 to perform axonal tracing and validated the presence of projections in our patch area (Figure 1A and 1B white arrows). Once we observed the presence of projections, we implanted the electrical stimulation electrode in S2, the optopatch pipette in dorsal striatum and an electrode in contralateral S2 in order to record LFP as shown on Figure 1C. Representative traces of patched MSN from dorsal striatum as well as S2 contralateral activity are shown on Figure 1D. In order to discriminate the dopamine receptor type present on our patched MSN, we pulsed blue light thanks to the optical fiber inserted in the patch pipette and recorded the MSN spontaneous activity (Figure 1E). Cells with D2 receptor are ChR2+ and respond to the light stimulation by depolarizing their membrane potential. On the contrary, cells with D1 are ChR2- and will not respond to the light stimulation. An example of D1 and D2 receptor patched cells filled with biocytin and confirmed with immunochemistry can be observed in Figure 1F. The intrinsic properties of recorded direct and indirect MSNs (D1 MSNs and D2 MSNs respectively) are presented on Table 6. We can observe a significant difference in the input resistance of dorsolateral direct and indirect MSNs, as previously described in mice (Reig and Silberberg, 2014; Ketzef et al., 2017b). Regarding to the post-synaptic potential (PSP) evoked during downstate (representatives traces Figure 1G and 1H), we observed a significant diminution of the mean amplitude after LY injection (Figure 1I) (control= 17.10 mV, LY= 11.69 mV, p=0.0005) linked to a significant diminution of the slope (Figure 1J) (control= 0.035, LY= 0.024, p=00024). This PSP amplitude diminution appears in both D1 MSNs and D2 MSNs and no significant differences is observed between both after LY injection (D1 MSNs= 62.54%, D2 MSNs= 71.61%) (Figure 1K). These results indicate that *in-vivo*, a LY i.p. injection affects the evoked glutamatergic response of MSNs. Indeed, the PSP amplitude as well as the slope are decreased.

**Table 6:**
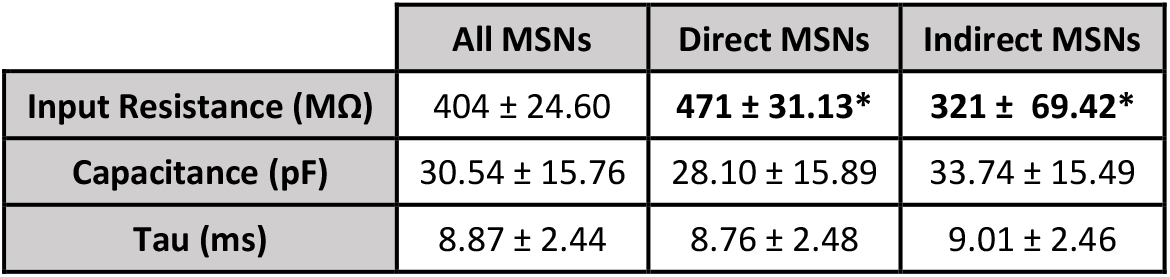
Intrinsic properties of recorded direct and indirect MSNs *in vivo*. All values are means ± standard deviation. n=38 (21 direct MSNs; 17 indirect MSNs). **\*** p<0.05, differences found in the input resistance when comparing direct and indirect MSNs.

|  | All MSNs | Direct MSNs | Indirect MSNs |
| --- | --- | --- | --- |
| **Input Resistance (MΩ)** | 404 ± 24.60 | **471 ± 31.13\*** | **321 ± 69.42\*** |
| **Capacitance (pF)** | 30.54 ± 15.76 | 28.10 ± 15.89 | 33.74 ± 15.49 |
| **Tau (ms)** | 8.87 ± 2.44 | 8.76 ± 2.48 | 9.01 ± 2.46 |

### CCK2R inhibition affects membrane active properties by reducing spike threshold and rheobase

MSNs are recorded in the dorsolateral region of the striatum (Figure 2A). After demonstrating that an injection of LY impacted direct and indirect pathway MSNs in the same direction (Figure 1K), *ex-vivo* recordings are performed without discriminating the type of dopaminergic receptors expressed by the MSNs. MSNs are identified thanks to their specific spiking pattern and I-V curve (Figure 2B and 2C). MSNs are hyperpolarized with an RMP between −70 mV and −88 mV, a K+ ramp that delayed the firing of the first spike and regular firing frequency. On the I-V curve we validate the presence of the inward rectification. As it is shown on Table 7, an addition of LY does not affect any passive membrane properties. On the contrary, regarding to the active membrane properties, (Table 8), LY affect the spike threshold and the rheobase. The mean spiking threshold presents a significant decrease after a LY infusion (control= −40.72 mV, LY= −49.02 mV, p=0.0313) (Figure 2D). Also, the mean rheobase presents a significant decrease after a LY infusion (control= 120 pA, LY= 68.33 pA, p=0.0313) (Figure 2E). A representative example of recorded spikes of the same cell before and after the infusion of LY (respectively control and LY) is shown on Figure 2F. Thus, these results indicate that CCK2R plays a role in the MSNs excitability.

**Table 7:** Membrane passive properties with or without CCK2R antagonist. All values are mean ± SEM. n=20. No significant resistance.

|  | Before LY | After LY | p value |
| --- | --- | --- | --- |
| **Ri (MΩ)** | 253.06 ± 52.76 | 220.21 ± 37.29 | ns |
| **Injected current (pA)** | -5.67 ± 11.72 | -11.18 ± 10.30 | ns |
| **Conductance (S)** | -0.33 ± 0.28 | -0.16 ± 0.16 | ns |
| **Membrane potential (mV)** | -76.95 ± 1.37 | -76.00 ± 1.70 | ns |

**Table 8:**
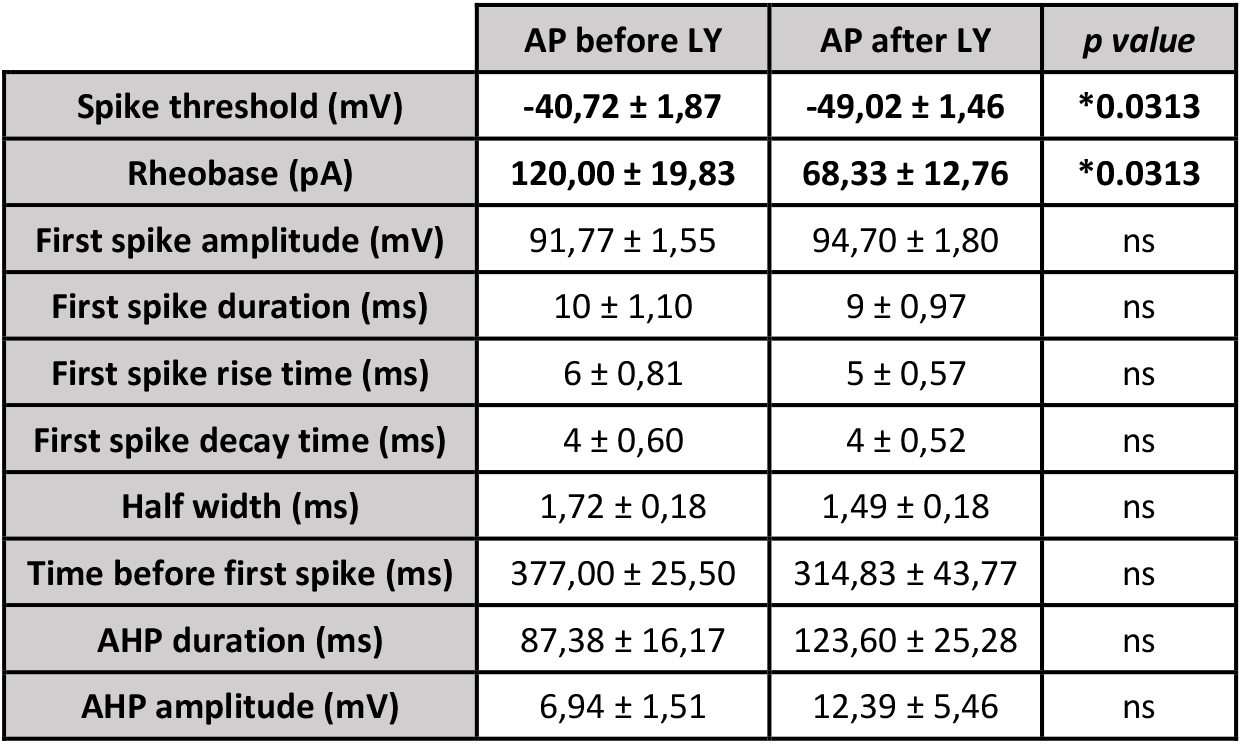
Membrane active properties with or without CCK2R antagonist. All values are mean ± SEM. n=6. *p<0.05, signed rank Wilcoxon test.

|  | AP before LY | AP after LY | p value |
| --- | --- | --- | --- |
| Spike threshold (mV) | -40,72 ± 1,87 | -49,02 ± 1,46 | *0.0313 |
| Rheobase (pA) | 120,00 ± 19,83 | 68,33 ± 12,76 | *0.0313 |
| First spike amplitude (mV) | 91,77 ± 1,55 | 94,70 ± 1,80 | ns |
| First spike duration (ms) | 10 ± 1,10 | 9 ± 0,97 | ns |
| First spike rise time (ms) | 6 ± 0,81 | 5 ± 0,57 | ns |
| First spike decay time (ms) | 4 ± 0,60 | 4 ± 0,52 | ns |
| Half width (ms) | 1,72 ± 0,18 | 1,49 ± 0,18 | ns |
| Time before first spike (ms) | 377,00 ± 25,50 | 314,83 ± 43,77 | ns |
| AHP duration (ms) | 87,38 ± 16,17 | 123,60 ± 25,28 | ns |
| AHP amplitude (mV) | 6,94 ± 1,51 | 12,39 ± 5,46 | ns |

### CCK2R inhibition affects the evoked glutamatergic response by reducing the EPSC amplitude *ex-vivo*

A stimulation electrode was placed in the layer V of somatosensorial cortex S2 (Figure 3A). An electrical stimulation was applied and the evoked current in the MSN was recorded in voltage-clamp (Figure 3B). The mean EPSC duration presented a significant decrease after a LY infusion (control= 45.13 ms, LY= 42.82 ms, p=0.0237) (Figure 3C). Moreover, the recordings showed a significative reduction of mean EPSC’s amplitude after LY infusion (Absolute mean: control= 166.02 mV, LY=153.40 mV, p=0.0120. Significative reduction of normalized mean: control= 100%, LY= 92.98%, p=0.0208) (Figure 3D). It is also illustrated by a representative EPSC from the same cell before and after LY infusion (Figure 3E). However, when applying Lorglumide (CCK1R antagonist) most recordings demonstrated a slight increase of mean EPSC’s amplitude (control= 116,46 mV, Lorglumide= 134,39 mV) (Supplementary data Figure 6B) and a significant decrease of mean EPSC duration (control= 43 ms, Lorglumide= 29,61 ms, p=0.0156) (supplementary data Figure 6A). Therefore, CCK2R plays also a role in the corticostriatal efficiency.

**Figure 6:**
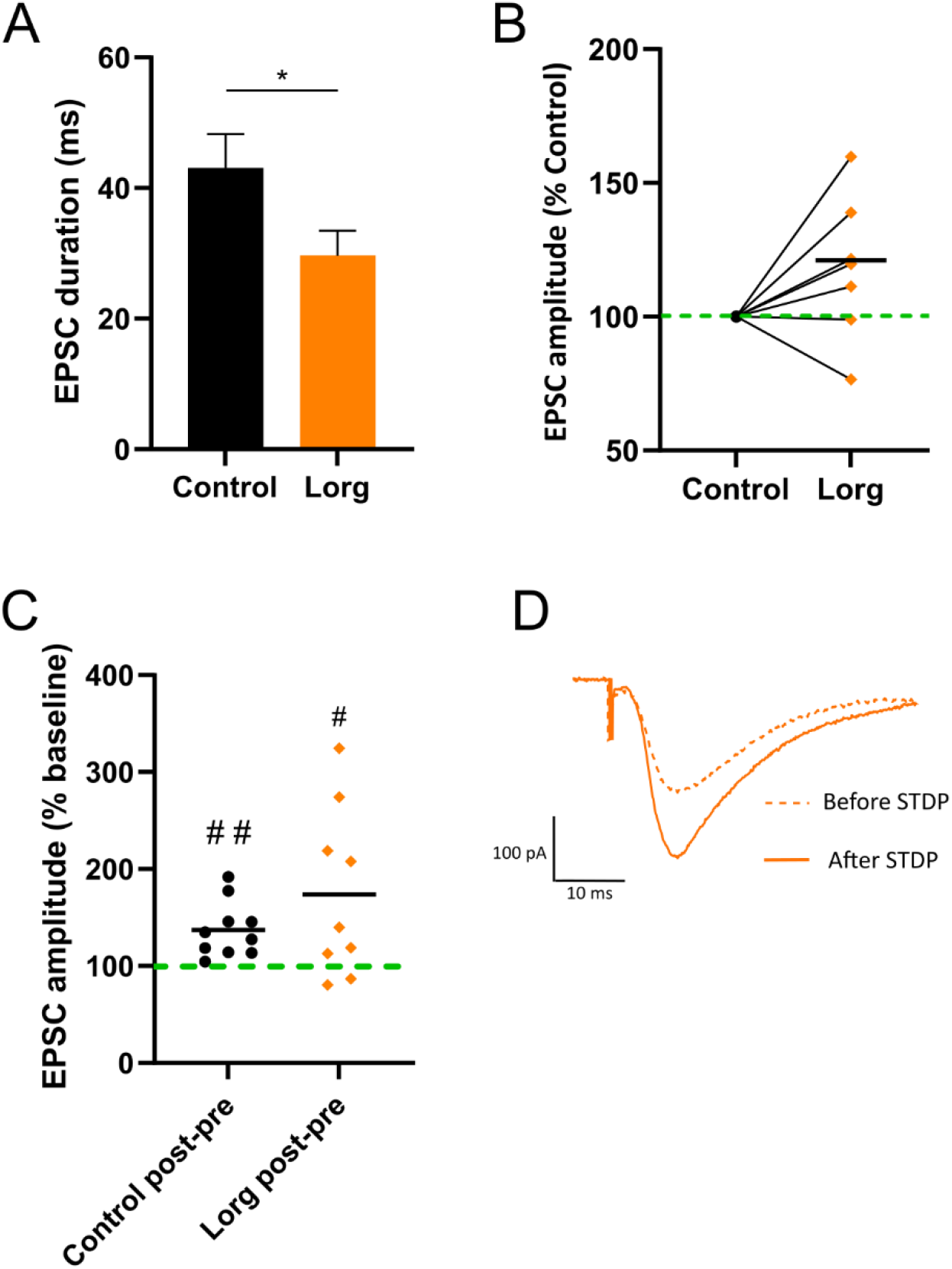
Lorglumide effects on the evoked glutamatergic response and STDP long term potentiation *ex-vivo*. **A** EPSC duration without and with Lorglumide, (*p= 0.0156, n= 7, Wilcoxon signed rank test). **B** EPSC amplitude of Lorglumide condition in % of the control condition (n=7). **C** Normalised EPSC amplitude in control condition and in Lorglumide condition, (control n=10 and Lorglumide n=9, #p=0.0340 Lorglumide vs baseline, one sample t test, ##p=0.0026 control vs baseline, one sample t test). **D** Evoked glutamatergic response from a representative recorded MSN in Lorglumide condition.

### CCK2R inhibition induces a switch on STDP Long Term Potentiation *ex-vivo*

To induce plasticity, we use a post-pre pairing where a MSN action potential is evoked few milliseconds before a cortical stimulation, 100 times at 1Hz (Figure 3F). In control conditions, the post-pre pairing induces a significant LTP (mean EPSC= 137.3% of baseline, p= 0.0026) whereas in LY condition this pairing induces a significant LTD (mean EPSC= 82.71% of baseline, p=0.0190) (Figure 3G). On the two representative recordings (Figure 3I), we show that the input resistance (Ri) changes were less than 20% between the base and the long term (cells with a Ri outside of this range are excluded of the analysis). This switch of plasticity in LY condition is not observed in the case of Lorglumide (CCK1R antagonist) addition in place of LY. Indeed, as in control condition, a post-pre pairing induces a significant LTP in Lorglumide condition (mean EPSC= 173.8% of baseline, p= 0.0340) (Supplementary data Figure 6C). These results suggest that CCK signaling is essential for LTP and the use of antagonist CCK2R induce a switch of plasticity. In control condition the LTP observed is presynaptic as indicated by the CV^2^ (Figure 3J). In LY condition, the LTD observed is postsynaptic. CV^2^ is calculated as described in <u>A Practical Guide to Using CV Analysis for Determining the Locus of Synaptic Plasticity</u> (*Brock JA and al., 2020*): (SD/μ)2, where μ is the mean.

### CCK2R inhibition impairs male motor behavior

#### LY and CCK8 effect on motor coordination

Male rats receiving LY injection show a reduced performance on Rotarod test compared to the control group (score of falling latency normalized to control group: LY= −16.73, p=0.0331) (Figure 4B) demonstrating a disability on motor coordination and the implication of CCK2R. No difference is observed between the performance of the CCK8 group and the control and LY group (score of falling latency normalized to control group: CCK8= −4.971) (Supplementary data Figure 7A). Regarding to female rats, no difference is observed between the LY group and the control groups (score of falling latency normalized to control group: LY= −0.08) (Figure 4C). No difference is observed between the performance of the CCK8 group and the control and LY group (score of falling latency normalized to control group: CCK8= 4) (Supplementary data Figure 7B). Our results suggest a sex-dependent response to LY since LY impairs only the male motor coordination.

**Figure 7:**
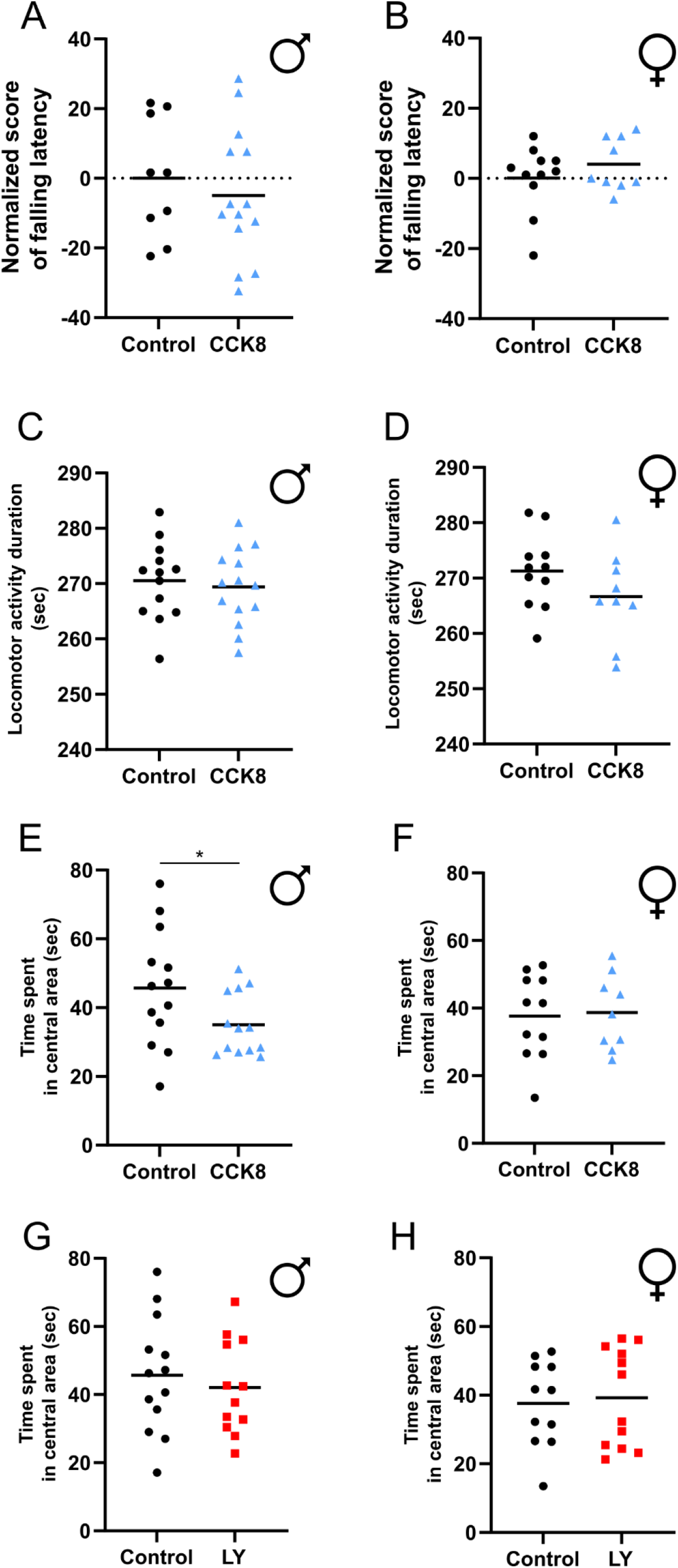
CCK8 effect on motor behavior. **A** Rotarod normalized score of falling latency (difference of falling latency between with and without injection of NaCl or CCK8, normalized to the value of the control group) for males’ rats (control n=12, CCK8 n=14). **B** Rotarod normalized score of falling latency for females’ rats in the Rotarod (control n=11, CCK8 n=9). **C** Males rats locomotor activity duration in control and CCK8 groups (control n=13, CCK8 n=14). **D** Female rats locomotor activity duration in control and CCK8 groups (control n=11, CCK8 n=9). **E** Time spent in central area by males’ rats’ control and CCK8 groups (*p=0.0453, n=13, Mann-Whitney test). **F** Time spent in central area by females rats control and CCK8 groups (control n=11, CCK8 n=9). **G** Time spent in central area by males’ rats control and LY groups (control n=13, LY n=12). **H** Time spent in central area by females rats control and LY groups (control n=11, LY n=12).

#### LY and CCK8 effect on behavior in the Open-field

Table 10 presents all parameters extracted from the OF test. We divide the OF in two areas: central (red square) and external (Figure 4A). We observe that male rats injected with LY have a reduced locomotor activity compared to the control group, by measuring the mean total movement duration in the whole OF (control= 270.50 sec, LY= 265.53 sec, p= 0.0457) (Figure 4D). The CCK8 group mean total movement duration did not differ from the control group (CCK8= 269.39 sec) (Supplementary data Figure 7C). Regarding to the total movement duration, females’ rats did not show differences between control and LY groups (control= 271.25 sec, LY= 266,56 sec) (4E). The CCK8 group total movement duration did not differ from the control group (CCK8= 266,63) (Supplementary data Figure 7D). Our results confirm our first observation with Rotarod: a sex-dependent response to LY since LY injection reduces only the male locomotor activity. In line with the known anxiogenic effect of CCK (Bradwejn et al., 1992; Daugé and Léna, 1998b; Wang et al., 2005), male rats injected with CCK8 show a reduced time spent in the central area of the OF (control= 45.68 sec, CCK8= 35.06 sec, p= 0.0453) (Supplementary data Figure 7E). This effect is not observed in the female’s rats (control= 37.64 sec, CCK8= 38.70 sec) (Supplementary data Figure 7F). In addition, LY injection did not modify the time spent in the central area for both sexes indicating an absence of anxiolytic effect when CCK2R is blocked (male LY= 42.13 sec, female LY= 39.20) (Supplementary data Figure 7G and 7H).

**Table 10:**
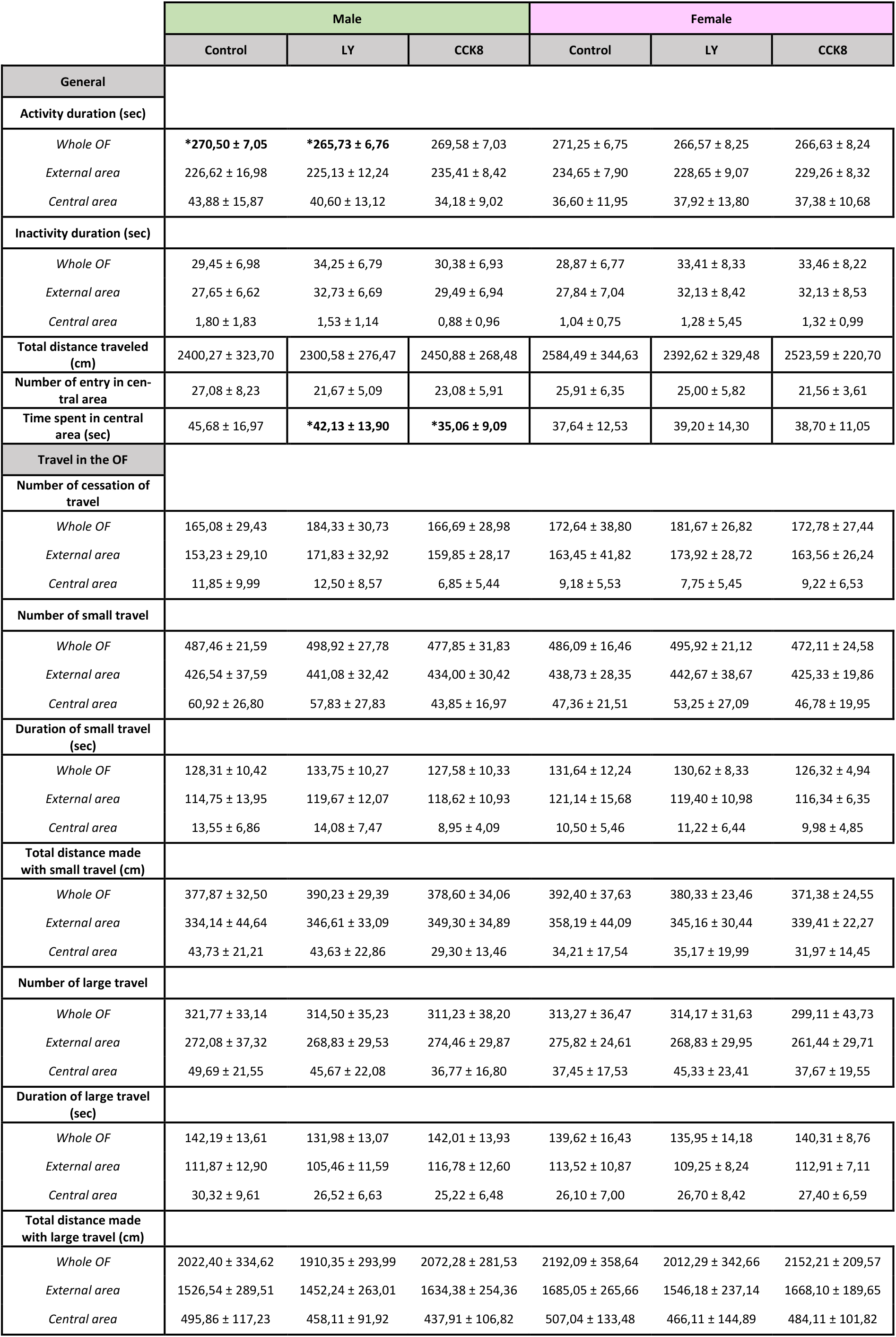
Extracted parameters from the Open-Field test. All values are mean and SD. For males, control, LY and CCK8 groups are respectively n=13, n= 12 and n=14. For females, control, LY and CCK8 groups are respectively n=11, n= 12 and n=9. * Significant differences are described in corresponding figure.

#### LY and CCK8 effect on postural recovery

The stepping test did not allow us to observe a difference in the postural recovery. Indeed, we did not see any effect of LY or CCK8 injection in the number of recoveries for any paw, in any of the sexes (Table 9).

**Table 9:**
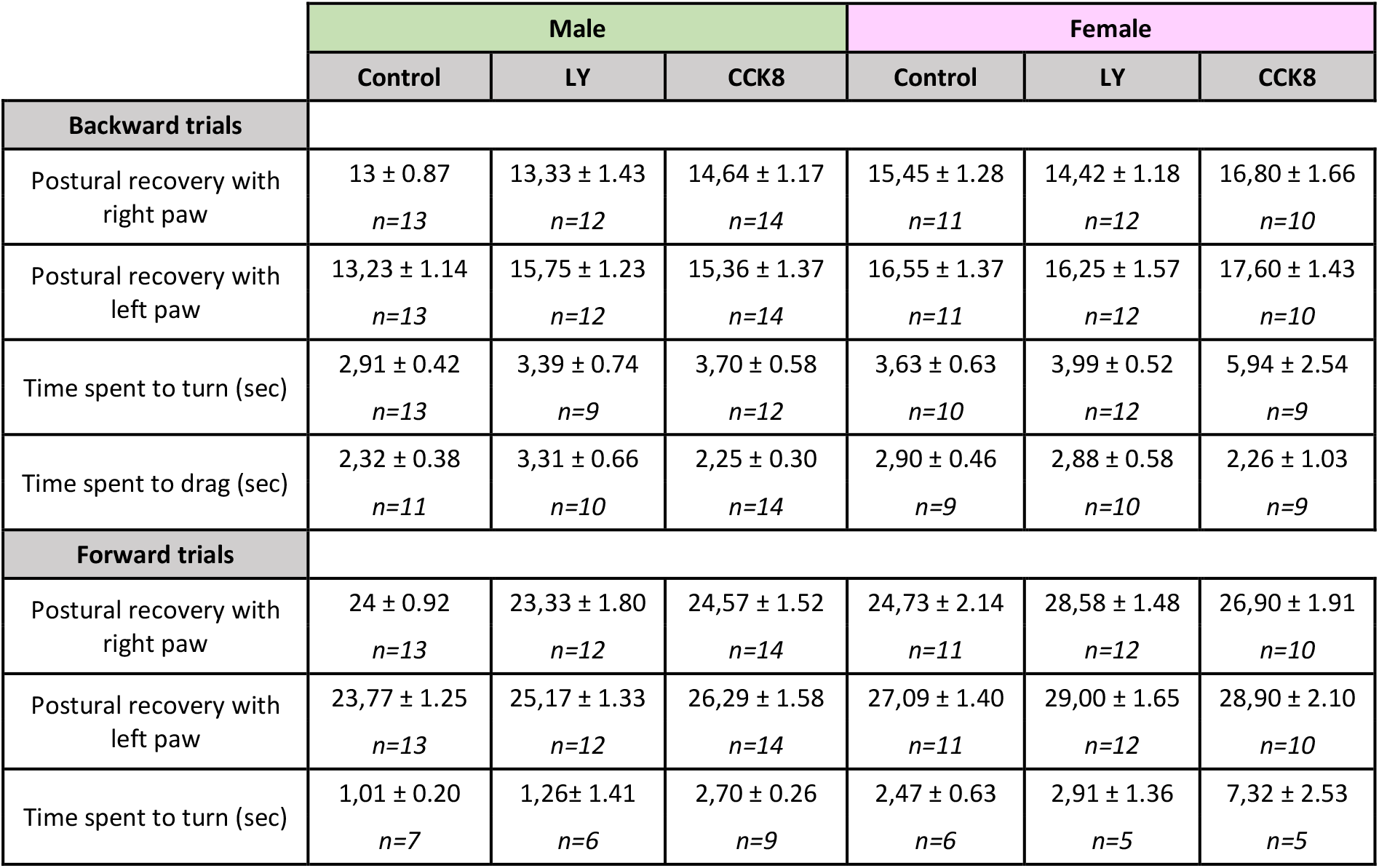
Evaluated characteristics with the stepping test. All values are mean per group ± SEM. n are indicated below the mean. No significant difference.

### CCK2R inhibition induces sex-dependent modification of c-Fos activity in the striatum

C-Fos positive cells density is measured as described on Figure 5A-C with an unbiased quantization technique to limit “over-counting”. This density is measured in each part of the striatum for males and females. In the male rats in each compartment there is a slight decrease in c-Fos labelling following LY injection (29.5 % of decrease for whole striatum). Due to the high degree of variability the quantification of the results is not significant (Figure 5D to 5H). Regarding the females, the c-Fos positive cells mean density is significantly higher in the ventro-medial striatum for the LY group compared to the control group (control= 75.73 mark/mm^2^, LY= 151.31 mark/mm2, p= 0.0221) (Figure 5M). This result is confirmed by the c-Fos positive cells density comparison in the whole striatum (control= 84.91 mark/mm2, LY= 129.42 mark/mm^2^) (Figure 5I). Our results suggest a sex-dependent effect of LY on c-Fos activity in the striatum.

**Figure 5:**
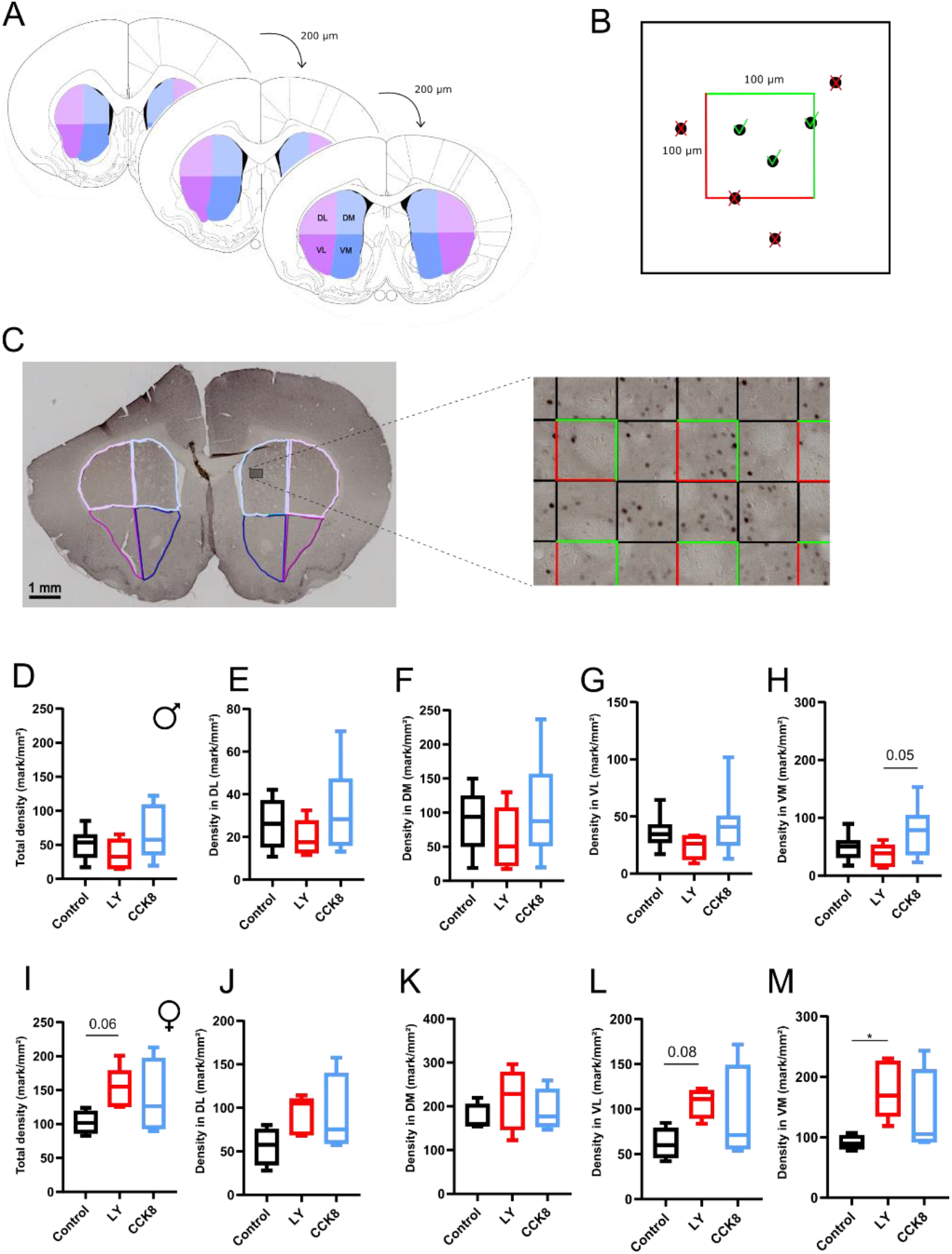
c-Fos immunolabelling analysis. **A** Free-floating serial coronal sections spaced of 200 μm were used for c-Fos immunolabelling. **B** Dissector system inclusion and exclusion was used to count c-Fos positive cells. **C** An example of a counting grid with dissector randomly positioned on the image. **D-H** c-Fos positive cells density (mark/mm^2^) in males’ striatum (control n=7, LY n=6, CCK8 n=8, mean on one hemisphere) D, E, F, G, H respectively whole striatum, DL, DM, VL, VM. In Figure H p= 0.05 LY vs CCK8, Kruskal-Wallis test following by Dunn’s multiple comparisons test. **I-M** c-Fos positive cells density (mark/mm^2^) in females’ striatum (control n=4, LY n=5, CCK8 n=4) I, J, K, L, M respectively whole striatum, DL, DM, VL, VM. In Figure I p=0.0586 control vs LY, Kruskal-Wallis test following by Dunn’s multiple comparisons test. In Figure L p=0.0792 control vs LY, Kruskal-Wallis test following by Dunn’s multiple comparisons test. In Figure M *p= 0.0221 control vs LY, Kruskal-Wallis test following by Dunn’s multiple comparisons test.

## Discussion

Neuropeptides, and especially CCK are known to modulate synaptic transmission in various brain structures, but how they control plasticity in corticostriatal glutamatergic transmission is not known. Our study is the first to demonstrate that CCK tightly controls STDP timing at the corticostriatal synapse. Using both *ex-vivo* and *in-vivo* electrophysiological approaches, we are able to demonstrate that the blockade of the specific receptor CCK2 strongly decreases the corticostriatal transmission. Furthermore, the *ex-vivo* approach using patch-clamp of striatal brain slices allows us to show that CCK is a key mediator of the synaptic plasticity induced by a STDP protocol. Indeed, when CCK2R is pharmacologically blocked, the synaptic plasticity is switched to a LTD in place of a LTP. Finally, we showed that CCK is involved in motor response via a sex-dependent mechanism that we do not elucidate yet. Indeed, blocking CCK2R impairs males motor coordination and reduces their general motor activity without significant alteration in female rats.

Interneurons expressing CCK are found in dorsal striatum and the presence of CCK is confirmed at mRNA and protein level (Muñoz-Manchado et al., 2018). These interneurons also co-express Vip (Takagi et al., 1984; Theriault and Landis, 1987; Hökfelt et al., 1988) or Th (Tepper et al., 2010). However, these CCK cells are sparse and Muñoz-Manchado et al. propose the hypothesis that these cells could be misguided CCK cells heading for cortical structures (Muñoz-Manchado et al., 2018). In the striatum, the low level of CCK-immunoreactive neuronal body cells and high level of CCK-like immunoreactivity in axons and terminals (Hökfelt et al., 1988) suggest an extrinsic origin of CCK. If the ventral striatum receive CCK+ projection from nucleus tractus solitarii (NTS)(Wang et al., 1992), the CCK present in the dorsal striatum originates from the cortex (Burgunder and Young III, 1990; You et al., 1994). Dorsal and ventral striatum also receive CCK from the mesencephalic region by dopaminergic fibers (Hökfelt et al., 1988). We should also note the existence of projections expressing CCK gene from the thalamus to the striatum (Burgunder and Young, 1988). The colocalization of CCK with neurotransmitters such as glutamate in the corticostriatal pathway (Bradwejn and Vasar, 2013) and dopamine from mesencephalon (Hökfelt et al., 1988) confirms the possible involvement of CCK in modulating transmission and synaptic plasticity. Then, given the many CCK fibers projection to the striatum from cortex and mesencephalon, as well as the presence of CCK+ interneurons in the striatum, CCK could directly modulate striatal inputs and information processing within the basal ganglia. Activation of CCK2R by CCK increases the activity of phospholipase C (PLC) which enhances the hydrolysis of phosphatidylinositol 4,5-biphosphate (PIP_2_) into inositol triphosphate (IP3) and diacylglycerol (DAG). IP3 increases intracellular Ca^2+^release when DAG activates protein kinase C (PKC) (Wank, 1995). So, a CCK2R activation will lead to an increase of intracellular Ca^2+^ and PKC activation. Further investigations are required to know where these receptors are localized at the synapse level. It could be on pre or post synaptic membrane, but we can also suggest that, as in (Crosby et al., 2018), CCK2R could be present on astrocytes involved in the synapse. In this case the activation of CCK2R could modulate synaptic processes via the astrocyte response.

### CCK2R antagonist changes corticostriatal synaptic transmission

Regarding our results on the corticostriatal synaptic transmission, we highlight in the *ex-vivo* electrophysiology recordings that blocking CCK2R with LY225910 reduces the amplitude of the evoked EPSC. In the *in-vivo* electrophysiology recordings this blocking reduces the amplitude of the EPSP. This diminution of EPSPs’ amplitude is not dependent of dopaminergic receptor (DR), D1 or D2, present on the MSN, as we show on Figure 1K. Because the administration of LY225910 was made by infusion in the extracellular bath in *ex-vivo* electrophysiology or by IP injection in *in-vivo* electrophysiology, the alteration of the transmission can be originated from the presynaptic part (cortical function) or from the postsynaptic (striatal function). In the cortex CCK is found in pyramidal neurons as well as in CCK-GABA interneurons (Taniguchi et al., 2011; Whissell et al., 2015), the latter being relatively more abundant in the secondary cortical area (as S2). Then, regarding our data, two hypotheses can be proposed. Firstly, the antagonist of CCK2R can act on the receptors present in the cortex, thus reducing the cortical neuronal activity and lowering the glutamate release. This reduction of activity will therefore have an effect on the MSN response. This hypothesis can be supported by some of our observations made on the *in-vivo* recordings where the cortical S2 LFP decreases after the LY injection (data not shown). Another hypothesis is that blocking CCK2R will stop the enhancement of glutamate release by CCK. Indeed, it is already shown that applying CCK8 or specific CCK2R agonist will enhance the EPSC amplitude by increasing the release of glutamate in the synapse (Migaud et al., 1994; Deng et al., 2010). This increase of glutamate release is dependent of the intracellular presynaptic Ca^2+^ release by PLC molecular pathway activation (i.e. the molecular pathway activated by CCK2R activation) and on the inhibition of presynaptic *I*_K_ (Migaud et al., 1994; Deng et al., 2010). The increase of intracellular presynaptic Ca^2+^ enhances the number of readily releasable vesicles and the probability of release, both parameters being Ca^2+^ dependent (Zucker and Regehr, 2002; Deng et al., 2010). These observations have been made in the hippocampus (Migaud et al., 1994; Deng et al., 2010), another region with a high level of CCK. Preliminary experiments performed in *ex-vivo* electrophysiology in our laboratory also suggest an increase of MSN EPSC after an addition of CCK8s (data not shown). However, further experiments, such as local CCK2R antagonist puffs at the vicinity of the recording neurons, are required to better understand the mechanisms of the CCK-induced plasticity shift.

### CCK2R antagonist switches the plasticity from LTP to LTD

To our knowledge, we are the first to show an involvement of CCK in the switch of corticostriatal synaptic plasticity. The corticostriatal synapse has a particular form of plasticity, called anti-Hebbian. In fact, differently from other brain structures such as the hippocampus or the cortex, if a postsynaptic action potential (AP) of MSN comes after a cortical stimulation (protocol pre-post) it induces an LTD; on the contrary a postsynaptic AP of MSN before a cortical stimulation (protocol post-pre) induces an LTP (Fino et al., 2005). Our study also shows the crucial involvement of CCK in the LTP induction by STDP protocol. Indeed, when CCK2R is blocked with LY225910, a post-pre protocol induces an LTD in place of an LTP. Such a switch in STDP plasticity from non-Hebbian to Hebbian plasticity was already observed in corticostriatal synapse in experiments where GABA_A_ receptors were blocked (Paille et al., 2013). So CCK signaling could also operate a non-Hebbian/Hebbian switch. We could not rule out that this switch is independent of GABA signaling. Our mean variance analysis suggests that the control LTP observed is originated from the pre-synaptic element, and that the LTD induced in LY condition is mainly originated from the post-synaptic element. These results are in line with classic non-Hebbian LTP and LTD previously described (Fino et al., 2005).

Interestingly, some other studies have shown the importance of CCK in the LTP. In 2019, Chen and al. showed in mice that CCK2R activation by CCK is essential for LTP induced by high frequency stimulation (HFS) in the neocortex and allows for the encoding of auditory associative memory. Indeed, the use of CCK2R antagonists will block neocortical LTP as well as induce a deficit in the association of two tones in a behavioral experiments (Chen et al., 2019). In this study they also showed that the NMDA receptors trigger the CCK release involved in the neocortical LTP induction. In 2018, Crosby and al. also showed the involvement of CCK and CCK2R in an HFS induced LTP of the GABA synapse in the dorsomedial hypothalamus (DMH)(Crosby et al., 2018). As in our study, the CCK signaling is able to shift the polarity of the plasticity: when the HFS protocol induces an LTD in control condition, the addition of CCK induces an LTP. This shift of polarity is dependent on the activity of CCK2R on astrocytes; the activation of CCK2R by CCK combined with the activation of mGluR5 by glutamate leads to the release of ATP by the astrocytes. ATP binds the P2X receptor on GABA terminals and causes a prolonged increase in GABA release (Crosby et al., 2018). In the striatal plasticity, dopamine receptors present on MSN play a key role (Shen et al., 2008). When the striatal dopaminergic level is low, as for example in Parkinson disease, both forms of plasticity are altered suggesting the crucial role of dopaminergic transmission in the corticostriatal plasticity induction (Calabresi et al., 2007). Different parameters are responsible for either LTP or LTD induction. The stimulation of D2, Cav1.3 channels or mGluR associated to a presynaptic activity are required to induce an LTD, whereas a presynaptic activity and stimulation of a D1 or adenosine receptor A2A lead to an LTP (Wickens, 2009). Endocannabinoid (eCB) CB1 receptors are also essential for LTD induction in this glutamatergic synapse by a retrograde signaling of eCB secreted by post-synaptic element (Gerdeman et al., 2002; Lovinger and Mathur, 2012; Xu et al., 2018). CCK can also induce a retrograde signaling as it was previously described (Crosby et al., 2015): post-synaptic CCK induces NO secretion which enhances the GABA secretion from the pre-synaptic element. We can then wonder if LTD-eCB dependent could be the counterpart of an LTP CCK dependent both working via retrograde signaling of eCB or NO. Thus, since several neurotransmitters and neuromodulators can modulate the plasticity, further experiments by pharmacological approaches should be performed to identify the process by which CCK control the corticostriatal LTP. It would be very interesting to check if LTD induced by LY225910 application is dependent on CB1 receptor activation. Especially as in behavioral study Beinfeld and Connoly showed that CB1 activation inhibits the CCK release in rat hippocampal slices (Beinfeld and Connolly, 2001). Thus, it would be worth investigating if it is also the case in the striatum and if the LTD induced by eCB is dependent or not of a CCKergic transmission reduction. Preliminary work in our laboratory showed that if the presence of LY225910 during post-pre protocols switches an LTP to an LTD, it does not reverse the plasticity if it is applied before a pre-post protocol (LTD induction).

### CCK2R antagonist impairs male motor coordination

Although we do not notice sex differences in the electrophysiology experiments we show that the impact of LY225910 on motricity is sex-dependent. Indeed, LY225910 induces motor impairment only in males. Hökfelt previously suggested an influence of CCK on motor behavior. Indeed, he described that an i.p. injection of CCK2R antagonist on naïve unhabituated rats induces a significant decrease in spontaneous motility (Hökfelt et al., 2002). Furthermore, in 2001, Daugé and al. showed that CCK2R deficient mice presented a significantly lower percentage of success in the Rotarod test, and that their score did not improve after three trials in comparison to the wild type mice (Daugé et al., 2001). These authors suggested an impairment in coordinated skills or in equilibration muscle strength and tonus. However, no sex-dependent effects were observed in the Rotarod test for these mice, but significant differences were observed in the Open-field test between males and females concerning the locomotor activity (Daugé et al., 2001). In 2001, Kõks and al. also showed in homozygous (−/−) and heterozygous (+/−) deficient CCK2R male mice an impairment of Rotarod test performance, stronger in homozygous, and a general reduction of locomotor activity (Kõks et al., 2001). They explained the alteration of the motor behavior by a change in the dopaminergic system since the sensitivity of presynaptic dopamine receptor was increased in both homozygous and heterozygous animals, as well as postsynaptic dopamine receptor in homozygous animal. Their hypothesis was reinforced by the study of Abramov et al. (Abramov et al., 2004) in which the deletion of CCK2R, either in homozygous or heterozygous male mice, increased the affinity of D2 in subcortical structures. That phenomenon was not observed in female where the CCK2R deletion leads to an increased affinity of 5-HT_2_ receptor in the frontal cortex, probably responsible for compensatory changes in female mice. Thus, our study confirms that inhibiting CCK2R leads to the impairment of motor coordination. In addition, we suggest that there is also an intersexual difference regarding the effect of a CCK2R blocking. A difference links to sex is also observed in our experiment on the striatal c-Fos expression. Indeed, a CCK2R inhibition is correlated with an increase of c-Fos expression in females but with a decrease in males. A sex difference to CCKergic signaling function and effect was already described few times and very recently. In fact, CCK appears to be involved in female’s reproductive behavior (Liu and Cao, 2022; Yin et al., 2022). A sex difference is also observed in CCK neuron activity and CCK receptor signaling in modulating drug intake and drug-seeking in rodent (Ma and Giardino, 2022). These data emphasize the need to examine sex and hormonal influences in CCK response and more generally in biology and health. We do not observe sex difference in our *in-vivo* electrophysiological recordings indicating that the sex difference observed is more complex and probably involves other brain areas than the specific corticostriatal synapse.

Our study highlights the findings that CCK signaling is crucial in corticostriatal LTP induction and male motor behavior. The interaction between striatal CCK, dopamine and/or eCB should be further investigated. Additionally, it will be interesting to determine the role of the systemic production of CCK versus the central production to understand if the neurobiological processes that we describe are only governed by CCK originating from the brain. To conclude, by studying the striatum, our data reinforce the crucial role of CCK signaling in controlling and modulating the synaptic plasticity in many brain regions. In light of these results, the place of CCK in cerebral pathologies (or brain disorders) must be revisited in order to consider new therapeutic approaches.

## Supplementary data

The authors declare no competing financial interests

